# Extensive incorporation, polarisation and improved maturation of transplanted human cones in a murine cone degeneration model

**DOI:** 10.1101/2021.08.26.457641

**Authors:** Sylvia J Gasparini, Karen Tessmer, Miriam Reh, Stephanie Wieneke, Madalena Carido, Manuela Völkner, Oliver Borsch, Anka Swiersy, Marta Zuzic, Olivier Goureau, Thomas Kurth, Volker Busskamp, Günther Zeck, Mike O Karl, Marius Ader

## Abstract

Once human photoreceptors die, they do not regenerate, thus photoreceptor transplantation has emerged as a potential treatment approach for blinding diseases. Improvements in transplant organization, donor cell maturation and synaptic connectivity to the host will be critical in advancing this technology to clinical practice. Unlike the unstructured grafts of prior cell suspension transplantations into end-stage degeneration models, we describe extensive incorporation of iPSC retinal organoid-derived human photoreceptors into mice with cone dysfunction. This incorporative phenotype was validated in both cone-only as well as pan-photoreceptor transplantations. Rather than forming a glial barrier, Müller cells extend throughout the graft, even forming a common outer limiting membrane. Donor-host interaction appears to promote polarisation as well as development of morphological features critical for light detection, namely formation of inner and well stacked outer segments oriented towards the RPE. Putative synapse formation and graft function is evident both at a structural and electrophysiological level. Overall, these results show that human photoreceptors interact readily with a partially degenerated retina. Moreover, incorporation into the host retina appears to be beneficial to graft maturation, polarisation and function.

**Highlights:**

- Generation of the first human iPSC cone reporter line
- Human cones extensively incorporate into the retina of mice with cone degeneration
- Donor cone age and time in vivo are important factors for transplant incorporation
- Incorporation into the host retina correlates with graft polarisation
- Improved photoreceptor maturation after transplantation in vivo vs. in vitro
- Re-establishment of cone-mediated light-responses in the cone deficient mouse

## Introduction

Vision impairment represents the most prevalent disability in the industrialized world and very few treatment options exist (1). Many blinding diseases are characterized by the progressive loss of photoreceptor cells which lack the ability to regenerate in mammals, including humans. Photoreceptor transplantation therapy has thus been proposed as a treatment modality in which healthy donor cells replace those that have been lost. Cell replacement therapies are an attractive option for retinal diseases – the eye is an organ which is self-contained and partially immune privileged, minimizing the risk of unwanted cell migration and rejection (2). Additionally, the eye is readily accessible and easily monitored. The cone rich foveal region, which is extremely important for human vision, facilitating tasks such as reading, facial recognition and driving, is relatively small, reducing the amount of donor cells required. Within the fovea there are only around 200 000 cones (3) – a number of cells which is readily produced with current organoid technology. However, to date no human cone-specific reporter line has been described and no efficient cell surface markers have been identified to facilitate effective sorting of donor cones. Although a marker panel for cone enrichment has been suggested, this provided low purity and yield (4).

Several recent studies have utilized human stem cell derived retinal organoids as a source of either photoreceptor or retinal sheets for transplantation, particularly in end stage-degeneration models. While some improvements in vision have been reported through the use of retinal sheets (5-9), these grafted sheets were largely disorganized, with extensive rosette formation and the complication of donor photoreceptors mostly synapsing to donor second order neurons rather than host cells. For human cone cell suspension transplantations, while functionality has recently been reported from two different research groups, grafts appeared disordered with little evidence of cell polarisation (10) (i.e. inner and outer segments oriented towards RPE, axons extended towards the second order neurons) or showed some polarisation but poor general transplant cell survival (11, 12).

The aforementioned studies mostly focused on transplantation into models of severe end-stage degeneration, particularly the rd1 mouse model, where no host photoreceptors remained. While these proof-of-concept studies are of utmost importance for the development of photoreceptor replacement therapies, the early onset and severity of the rd1 phenotype represents a rather atypical pathology in regard to retinal degeneration patients. Complete photoreceptor cell loss only occurs in patients in very late stages of retinal disease, while in AMD, the most prevalent retinal degenerative disease, massive photoreceptor degeneration, called geographic atrophy (GA), is locally restricted. Additionally, a highly degenerated environment may not be conducive to graft survival, organized graft integration or synaptic connectivity with the host retina. In humans, extensive glial scarring and neural retinal remodeling may render end stage transplantations challenging (13). It is not yet known in which retinal disease type or at what degenerative stage photoreceptor replacement therapies would be most effective. Here, we therefore used the cone photoreceptor function loss mouse (Cpfl1), in which cones are dysfunctional and rapidly degenerate while rods remain largely unaffected (14), in order to determine whether human cones can integrate into the existing host photoreceptor layer.

In this study, a cone-specific human iPSC GFP-reporter line was generated in order to facilitate FAC sorting of an enriched cone population from retinal organoids. We used an optimized differentiation protocol which generates cone-rich retinal organoids ensuring a large population of transplantable cone cells. We aimed to investigate how transplanted cones mature, as well as how the donor-host interaction changes over time after transplantation. Results show long-term survival for up to six months in mouse recipients, extensive and polarised incorporation into the remaining mouse outer nuclear (photoreceptor) layer and interaction with host Müller glia and second order neurons. Human graft incorporation was further validated through the use of donor photoreceptors from a pan-photoreceptor reporter iPSC line. Moreover, photoreceptor graft maturation and polarisation was enhanced by donor-host interaction, as shown by histology, ultrastructural analysis and transcriptomics. Human photoreceptor transplants ultimately led to the re-establishment of cone-mediated light-responses in the cone deficient mouse.

## Results

### Validation of a human cone reporter iPSC-line to produce a transplantable population of human cones

A human iPSC line carrying GFP under the control of the cone-specific mouse cone arrestin (mCar) promoter was generated using a piggyBac transposon system (mCar-GFP line). This did not affect karyotype (Fig S1B,C). Human mCar-GFP retinal organoids were produced using a modified version of a previously published protocol which has been shown to generate robust numbers of cone photoreceptors (Fig S1A)(15-17). The mCar-driven GFP signal was predominantly located in the outer neuroepithelial layer as to be expected for cones (Fig 1A). Reporter expression co-localised with human cone arrestin (ARR3) antibody staining and all ARR3 positive cells appeared to be GFP^+^, indicating the specificity and efficiency of this 5 reporter. The GFP^+^ cells were positive for the photoreceptor-specific markers CRX and recoverin, and also expressed more mature cone markers such as L/M Opsin, and S-opsin at day (D) 240 of in vitro differentiation (Fig 1B). Note that there are far more L/M opsin cones present in the organoids than S-opsin cones, as previously described (15). Markers of other retinal cell types, namely, rods (Nrl and Rhodopsin), Müller glia (Sox2 and GLAST/CRALBP), bipolar (PKC*α*) and amacrine/ganglion cells (HuC/D) did not colocalize with GFP (Fig S1D-F). For a more in-depth analysis of the cell identity of GFP expressing cells, next generation sequencing of FAC-sorted GFP^+^ and GFP^−^ cells was performed with D200, D270 and D370 retinal organoids. This analysis confirmed that GFP^+^ cells highly express cone specific genes such as ARR3, CNGB3, PDE6C and L and M opsins, whereas the negative fraction showed high expression of typical marker genes of other retinal cell-types including rods, Müller glia, bipolar cells and retinal ganglion cells (Fig 1B). Additionally, Gene ontology term analysis of differentially expressed genes in cones from D200 versus D270 organoids revealed an enrichment of cellular compartment pathways critical to photoreceptor function in D270 cones, indicating that D200 cones are not yet fully mature and undergo extensive molecular changes in the following 10 weeks (Fig 1C).

**Figure 1:**
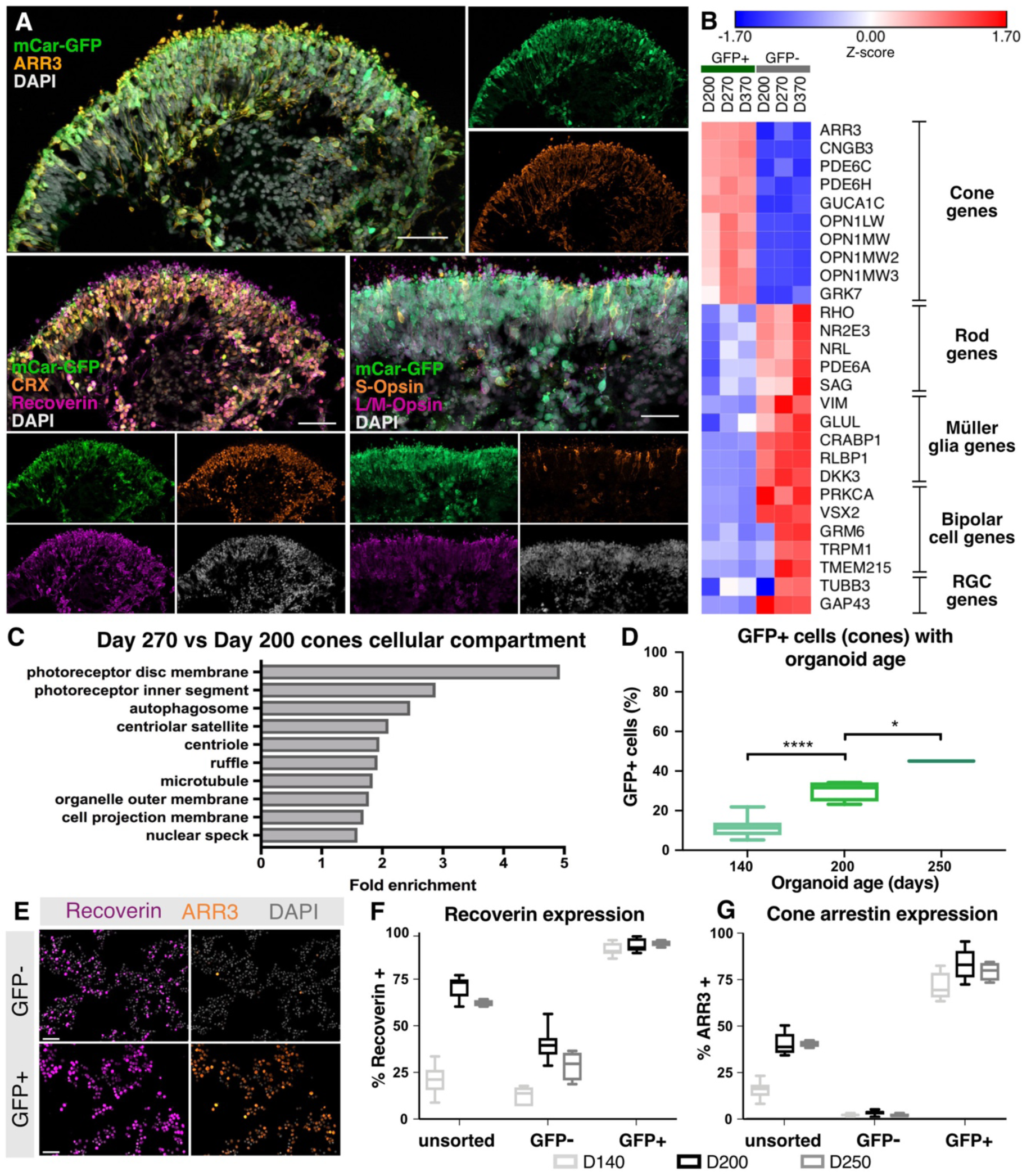
Generation and characterisation of a cone specific reporter line – D240 mCar-GFP derived retinal organoid cryosections show (A) co-staining of mCar-driven GFP with cone (ARR3, S-Opsin, L/M-opsin) and photoreceptor specific (CRX, recoverin) proteins. (B) Heat map of z-scores in major retinal cell type marker gene expression in GFP+ and GFP-cells sorted from mCar-GFP reporter organoids at D200, D270 and D370 post differentiation. (C) Gene ontology term cellular compartment over-representation analysis of D270 GFP+ cells compared with D200 GFP+ cells. (D) Proportion of GFP+ cells with organoid age. (E) Immunocytochemistry of GFP, recoverin and ARR3 expression in GFP+ and GFP-FAC-sorted fractions and quantification of immunocytochemical staining of (F) recoverin and (G) cone arrestin in unsorted, GFP+ and GFP-sorted fractions. Scale bars in all immunohistochemical images 50 μm.

To assess the proportion of cones in the organoids and the efficiency of reporter expression, FAC-sorting followed by immunocytochemical analysis was performed (Fig 1D-G). As expected, there was a significant increase in the proportion of GFP^+^ cells with organoid age (i.e. at D140, D200, D250), with up to 45% of cells determined to be GFP^+^ by D250 (Fig 1D). The FAC-sorted GFP^+^ fraction was found to be highly enriched in recoverin and ARR3 positive cells (Fig 1E-G), whereas the GFP^−^ fraction was almost entirely depleted of cone arrestin positive cells at all time points investigated (Fig 1G). This indicates that almost all cones are captured using the mCar-GFP reporter-based sorting system. With the confirmation of the cone identity of GFP^+^ cells, cones from D200 organoids were determined to be the most suitable population to perform transplantation studies, due to the robust number of relatively mature cone cells present, combined with a high degree of viability following dissociation and FACS purification. A smaller transplantation study using cones from D250 organoids was also performed for comparison.

### Human cones incorporate extensively into the host retina with longer post-transplantation times

Human cones were transplanted into the subretinal space (between the RPE and photoreceptor layer) of Cpfl1 mice, which received monthly vitreal triamcinolone acetonide injections for immune suppression from the time of transplantation. All transplanted cells expressed human ARR3 across the study timeframe and minimal immune reactivity of the host was observed (Fig S2A,B). Three weeks after transplantation, clusters of human cones survived in the subretinal space but did not interact extensively with the host outer nuclear layer (ONL). Donor cell clusters appeared mostly separated from the host ONL with few contact points (Fig 2A). Strikingly, 10 weeks after transplantation, large clusters (up to 30,000 μm^2^ per retinal section) of human cones were found to be partially incorporated into the Cpfl1 host ONL (Fig 2B) and appeared to incorporate further by 26 weeks (Fig 2C). Note that this phenomenon is not due to material transfer, which is frequently observed in mouse-to-mouse photoreceptor transplantations (18-20). Here, GFP^+^ cells are identifiable as human by human mitochondrial and human ARR3 expression, as well as through significantly larger and less dense nuclei than mouse photoreceptors (Fig S2, see also (10, 21)). Additionally, transcriptome analysis by next generation sequencing revealed the human origin of GFP^+^ cells isolated from transplanted retinas (see below).

**Figure 2:**
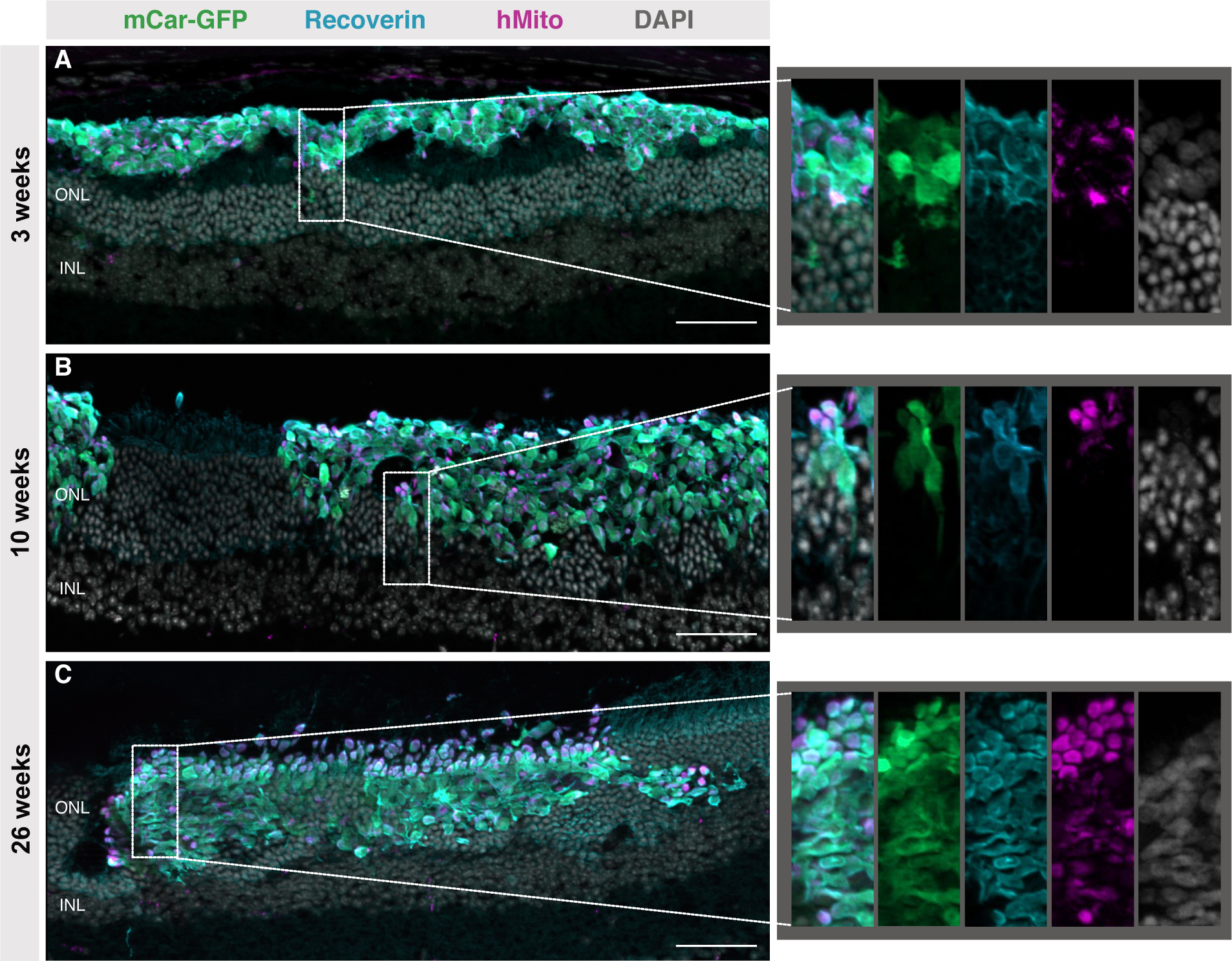
Extensive incorporation of transplanted cones into the Cpfl1 host retina with increased time since transplantation – Cryosections of retina transplanted with mCar-GFP^+^ cells at D200 post-differentiation stained with GFP, recoverin, human mitochondria and DAPI show (A) minimal donor-host interaction 3 weeks post-transplantation and (B) large cell clusters incorporated into the host retina at 10 weeks post-transplantation, with areas of round mitochondria rich outgrowths towards the RPE and axon like extensions projected towards the inner nuclear layer (see zoomed area). (C) By 26 weeks, grafts displayed even more abundant mitochondria rich outgrowths (see zoomed area). Scale bars in all immunohistochemical images 50 μm. RPE: retinal pigment epithelium, DAPI: 4ʹ,6-diamidino-2-phenylindole

### Maturation of human cones within Cpfl1 hosts

In addition to incorporating into the host ONL over time, human cones also appear to further mature in vivo. While at 3 weeks post-transplantation the donor cell mass was largely amorphous, by 10 weeks transplanted cones developed axon-like projections towards the host INL and mitochondria rich bulbous outgrowths towards the RPE. As photoreceptors are characterised by two distinctive compartments - namely the highly metabolic inner segment containing densely packed mitochondria and the unique light detecting outer segment, an elaborated primary cilium comprised of stacked disc membranes - the observed mitochondria rich bulbous outgrowths are indicative of inner segment development (Fig 2A,B). These presumed inner segments were even more widespread by 26 weeks post-transplantation (Fig 2C). To confirm the inner segment identity of the bulbous mitochondria-rich outgrowths, retinal sections were stained with markers associated with inner and outer segments. Accordingly, PNA which is specific for cone inner and outer segments, was bound in a non-localised fashion throughout the graft at 3 weeks. By 10 and even more prominently by 26 weeks, the PNA label was increasingly concentrated in mitochondria rich regions, i.e. the RPE facing edge of incorporated grafts and the rosette-like structures which occurred in some areas where mouse photoreceptors remained underneath the incorporating graft (Fig 3A). Peripherin-2 (PRPH2) staining of outer segments was not evident in the human cones at 3 weeks and only occasionally at 10 weeks post transplantation, however, by 26 weeks, PRPH2 was expressed in close association with the putative inner segments (hMito), also suggesting outer segment formation by this timepoint (Fig 3B).

**Figure 3:**
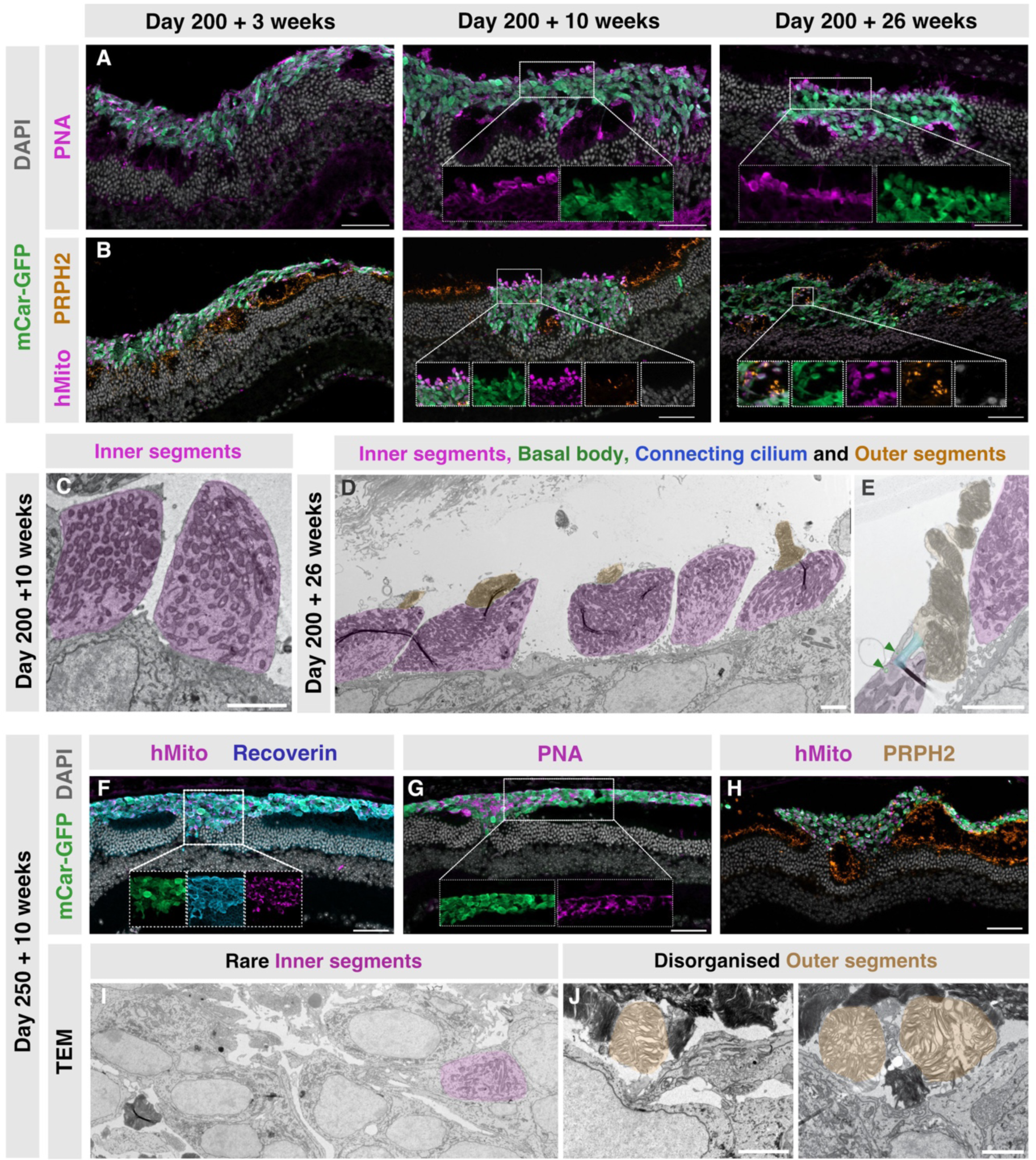
Graft development, polarisation and inner and outer segment formation – Cryosections of retina transplanted with D200 mCar-GFP^+^ cells were stained with (A) PNA showing more localised PNA binding with longer transplantations times. (B) PRPH2 shows most abundant staining at 26 weeks post-transplantation. TEM of ultrathin sections of eyes transplanted with D200 cones revealed (C) inner segments (purple) at 10 weeks post transplantation, (D) inner (purple) and outer segments (orange) and (E) occasionally basal bodies (green arrows) and connecting cilium (blue overlay) at 26 weeks post-transplantation. Cryosections of retina transplanted with mCar-GFP^+^ cells at day 250 post-differentiation showed (F) minimal donor-host interaction and few mitochondria rich outgrowths, (G) dispersed PNA binding and (H) little PRPH2 staining. TEM of the D250 transplanted cones showed (I) few inner segments and (J) occasional disorganised outer segments. Scale bars in all immunohistochemical images 50 μm and for all TEM images 2 μm. IS: Inner segment, OS: Outer segment, BB: Basal body, CC: Connecting cilium, TEM: Transmission electron microscopy, PNA: Peanut agglutinin, PRPH2; Peripherin2

To investigate the extent of photoreceptor maturation further, grafts were examined at the ultrastructural level. Indeed, many examples of inner segments were seen at 10 weeks post-transplantation, whereas outer segments were not found (Fig 3C). By 26 weeks, however, numerous cones formed relatively well organized and tightly stacked outer segment-like structures which were sometimes found to be joined to inner segments via a connecting cilium, additionally identifiable by the characteristic basal bodies (Fig 3D,E). The cells displaying these photoreceptor specific features were confirmed to be of human origin by the distinctive size and morphology of the human cone nuclei (these are much larger and less electron dense than the mouse photoreceptor cells – e.g. Fig 5A,B) as well as through immunogold labelling of human specific ARR3 (Fig S2C).

**Figure 4:**
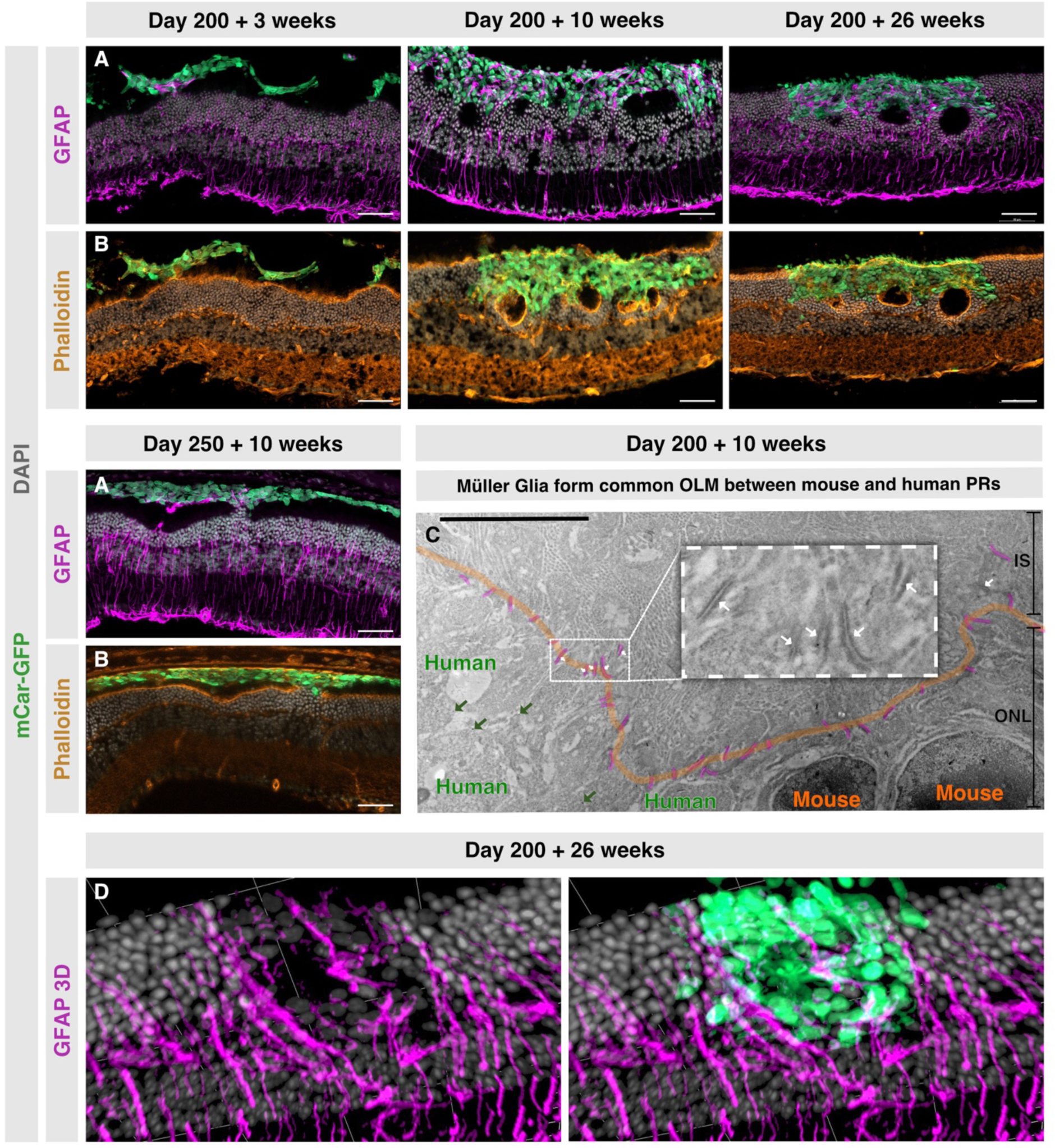
Host müller glia interaction with human cone grafts – Cryosections of retina transplanted with mCar-GFP+ cells at D200 or D250 post-differentiation showed (A) Müller glia beginning to extend processes into areas where the graft contacted host ONL (D200+3 weeks, D250+10 weeks), and extensively intermingling with grafts (D200+10 and D200+26 weeks) which had incorporated into the host ONL. (B) Phalloidin staining indicates that a common continuous OLM forms when the human cones incorporate into the host ONL. (C) Immunogold labelling confirms the formation of a common OLM between mouse and human photoreceptors. Dark green arrows indicate examples of immunogold 10nm labelling of human ARR3, thick orange line indicates the position of the OLM, yellow strokes indicate adherens junctions between mouse Müller glia and both mouse and human photoreceptors. (D) 3D reconstruction of GFAP positive Müller glia processes extending around human cones. Scale bars in all immunohistochemical images 50 μm, for CLEM images 6 μm and in 3D reconstruction grid lines are 50 μm. OLM: outer limiting membrane

**Figure 5:**
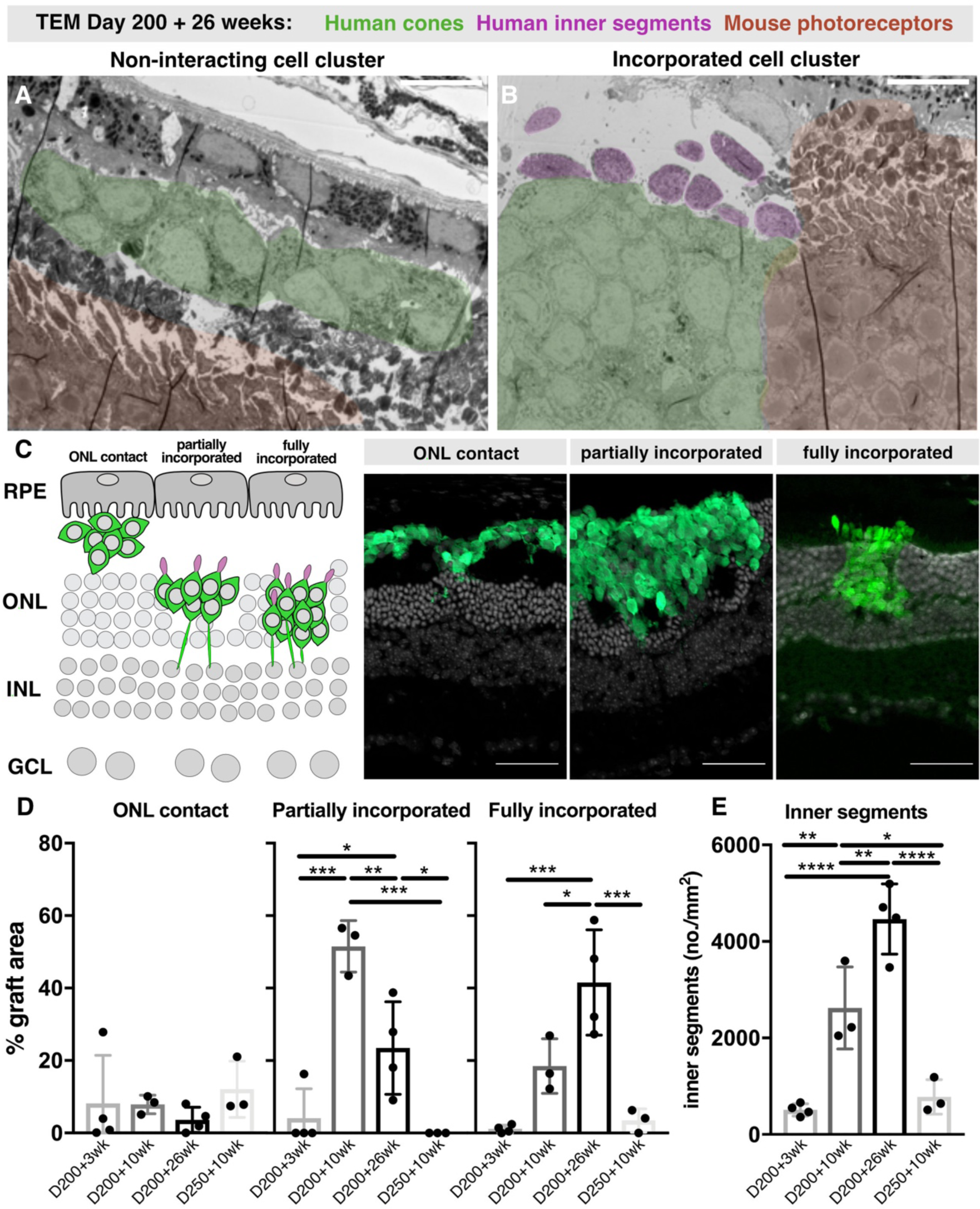
Interactive grafts more readily develop inner segments – Representative TEM of ultrathin retinal sections where some cone clusters (green overlay) within the same mouse eye (A) remain in the subretinal space or (B) incorporate into the host ONL (mouse photoreceptors orange overlay) and develop inner segments (purple overlay). (C) Schematic representation and example retinal cryosections for the classification of donor-host interaction into ONL contact, partially or fully incorporated. (D) Quantification of retinal cluster interaction with the host retina by area (n=3-4 eyes). (E) Number of mitochondria rich presumed IS at each timepoint (n=3-4 eyes). Scale bars in all immunohistochemical images 50 μm and for all TEM images 10 μm. IS: Inner segment, ONL: Outer nuclear layer, TEM: Transmission electron microscopy. Data displayed as mean and SD *p<0.05, **p≤0.01, ***p≤0.001, ****p≤0.0001

As inner and particularly outer segments took a long time to develop post-transplantation, we postulated that transplanting cones derived from older organoids might reduce the time required for the in vivo development of such mature photoreceptor specific features. Cones isolated from D250 retinal organoids were transplanted and assessed 10 weeks post-transplantation. Interestingly, unlike D200 cones, after 10 weeks in vivo most of the D250 grafts remained in the subretinal space, indicating a reduced capacity of the older cells to incorporate into the host ONL (Fig 3F). Much like D200 + 3 week transplantations, the D250 + 10 week grafts were presenting as a largely amorphous cell mass with few mitochondria rich or PRPH2 outgrowths evident and PNA label dispersed through the cell mass, rather than accumulating towards the RPE (Fig 3F,G,H). At an ultrastructural level, occasional inner segments as well as some outer segments were observed, however, the outer segments were highly disorganized and not tightly stacked (Fig 3I,J). Although photoreceptors of D250 + 10 week grafts (i.e. post-differentiation D320) are in total older than D200 + 10 week grafts (post-differentiation D270), they in comparison show a decreased capacity for incorporation and maturation. This suggests D200 cones are a preferable donor cell age. Together, these observations indicate that donor cone age and time in vivo are important factors for transplant incorporation and maturation.

### Müller glia incorporate transplanted cones into the host retina, forming a common OLM

In normal retinal physiology, photoreceptors are intermingled in a dense network of Müller glia processes that support photoreceptor structure, homeostasis, and function – even participating in the cone visual cycle (22). Therefore, the interaction between transplanted human cones and recipient Müller glia was assessed.

Immunohistochemical staining for GFAP revealed that in the D200 + 3 week and D250 + 10 week transplants, Müller glia processes extend into the graft only in limited areas where donor clusters start to make contact with the ONL, while no GFAP staining was observed within subretinal-located grafts (Fig 4A). By D200 + 10 weeks, rather than forming a glial barrier, Müller glia processes permeate throughout the graft (Fig 4A), and seemingly create an outer limiting membrane (OLM) in between the human nuclei and the subretinal space. Phalloidin staining supports this finding, showing an actin-dense band above the human nuclei which is continuous and in line with the host OLM, incorporating the clusters of human cones rather than excluding the xenogeneic cells (Fig 4B). This interaction was maintained at 26 weeks (Fig 4A, B, D).

These observations were confirmed by EM, where close association of Müller glia processes and human cones was evident. The continuous band of adherens junctions formed between them at the base of the inner segments strongly supports the OLM phenotype and resembles normal OLM structure. Furthermore, EM analysis again showed the continuity of the OLM between human cones and endogenous photoreceptors (Fig 4C).

Importantly, we also observed via EM that even within the same eye, it was primarily in these clusters of incorporated human cones that mature photoreceptor-specific features of inner and outer segments developed, whereas those clusters of cones which remained isolated in the subretinal space, without obvious interaction with host Müller glial processes, persisted largely amorphous (Fig 5A,B).

To quantify the extent of donor-host interaction at different experimental timepoints, total transplanted cell area was determined and the percentage of interacting grafts was calculated. Here, “ONL contact” was defined as areas where the cell mass remains in the sub-retinal-space but had points of contact with the ONL (Fig 5C). “Partially incorporated” graft was defined as areas where the transplant was in line with the host ONL (Fig 5C), but where some host photoreceptors remained beneath the graft and often formed small rosette-like structures. The graft was only considered “fully incorporated” when the transplant area appeared to replace stretches of host ONL, with direct contact to the INL and no rosettes or gaps evident (Fig 5C). Over half of the D200 + 10 week transplant cell clusters partially incorporated and a further 20% fully incorporated into the host ONL. By D200 + 26 weeks, over 40% of the graft area was fully incorporated. Both the D200 + 3 week and the D250 + 10 week samples only minimally interacted with the host retina (∼85% graft area non-interacting) (Fig 5D). Accordingly, only D200 + 10 week and D200 + 26 weeks grafts exhibited numerous mitochondrial rich outgrowths, i.e. inner segments (Fig 5E). If this were simply a factor of cell age, one would expect D250 + 10 week to display at least as many inner segments as D200 + 10 week grafts, however, in line with our previous observations, these only developed very few mitochondria-rich inner segments. Moreover, where inner segments did develop, these appear almost exclusively in areas where the host retina is directly contacted by the graft (Fig 3F), indicating that interaction with the host influences the maturation and development of photoreceptor-specific morphological features like inner segments.

### Cones mature more extensively in the mouse retinal environment compared to those maintained in retinal organoids in vitro

In order to further investigate whether the maturation trajectory of the retinal organoid-derived cones was influenced, as we suggest, by the host retinal environment, we compared the transcriptional profile of transplanted cones with cones from age-matched retinal organoids. D200 organoids were either maintained for a further 10 or 26 weeks (henceforth referred to as in vitro) or whole eye cups were dissociated at 10- and 26-weeks post transplantation (hereafter referred to as in vivo), and GFP^+^ cells recollected via FACS for RNA sequencing (Fig 6A). Interestingly, PCA analysis of the top 500 differentially regulated genes revealed that the greatest source of variance in the data separated clusters not depending on their age (D200 + 10 week and D200 + 26 week in vitro samples cluster closely together in PC1), but according to the time in vivo, indicating that maturation within the host retina indeed plays an important role (Fig 6B). More detailed gene overrepresentation analysis showed that molecular mechanisms, biological processes and cellular compartment pathways involved in light perception were highly and significantly enriched in the in vivo matured cone 12 samples (Fig 6C). Both L and M opsins as well as other outer segment related genes were highly upregulated in the in vivo matured samples – particularly after 26 weeks (Fig 6D). In the D200 + 26wk in vivo cones there was additionally enrichment in many mitochondrial and respiratory pathways compared to age-matched in vitro matured cones, indicating higher metabolic capacity in the in vivo matured cones (Fig 6 C, E). This analysis supports the histological and ultrastructural evidence that the host retinal environment promotes maturation of organoid-derived human cones, leading to enhanced inner and outer segment formation, which is critical to light detection.

**Figure 6:**
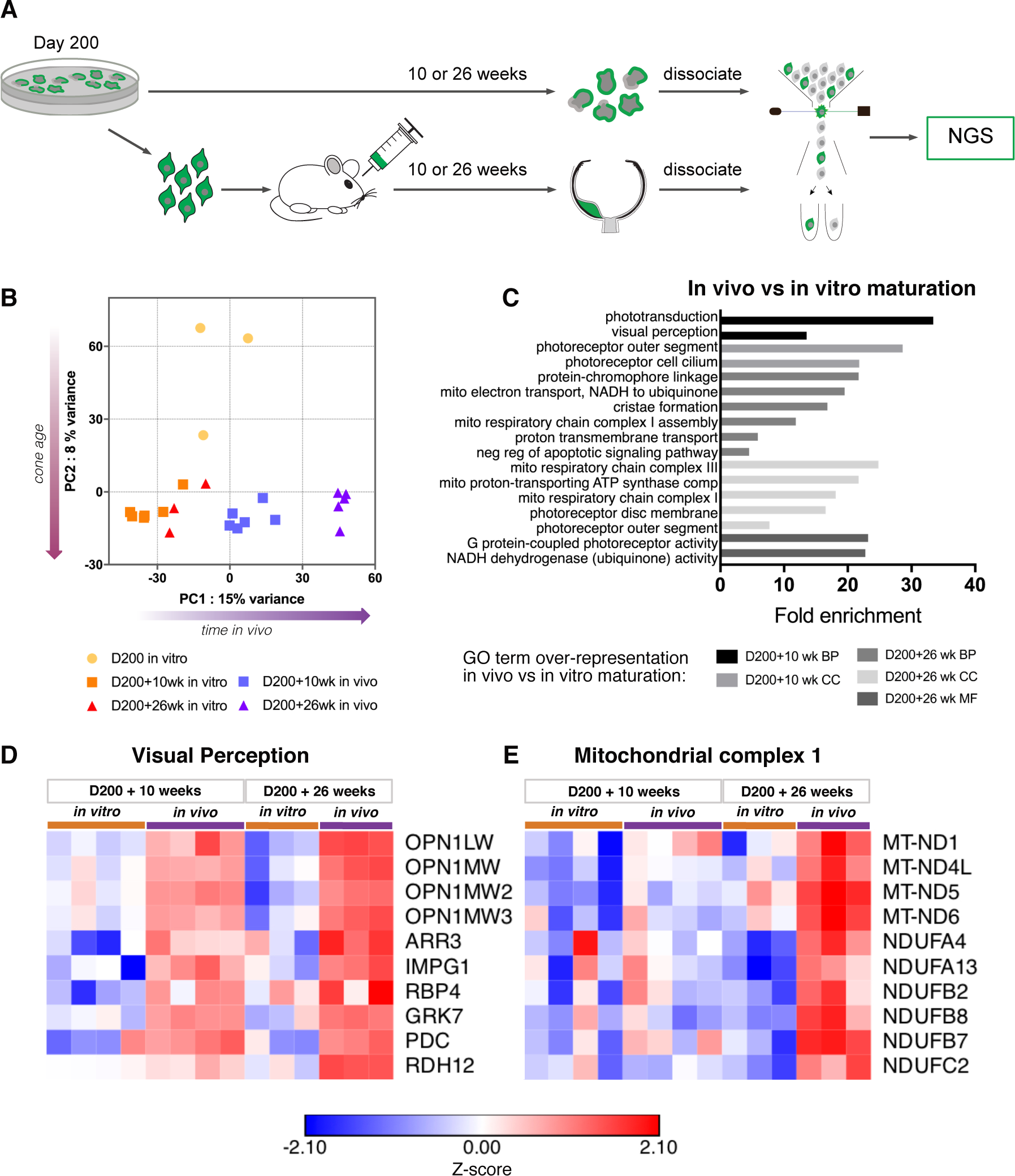
Transcriptional profiling of transplanted cones compared to age-matched organoid derived cones – (A) Schematic representation of mCar-GFP^+^ cone sequencing work-flow. (B) principal component analysis of the top 500 differentially regulated genes. (C) GO term pathway over representation analysis of in vivo matured vs in vitro matured cones. Heat maps of z-scores for genes involved in (D) visual perception and (E) mitochondrial complex 1.

### Validation of donor cell incorporation using the CRX iPSC cell line

To examine whether the incorporating capacity displayed by the human cones was specific to this cell line, we generated and transplanted photoreceptors from a CRX driven mCherry reporter iPSC line (23). CRX is expressed in retinal progenitors, rods and cones, with Crx-mCherry thus marking both photoreceptor cell types (Fig S3). FAC-sorted D200 Crx-mCherry^+^ cells were transplanted into Cpfl1 mice as per mCar-GFP^+^ cones. A remarkably similar phenotype was observed where Crx-mCherry^+^ photoreceptor transplants appeared to replace whole sections of mouse ONL (Fig 7A,B), with apical oriented inner segments (Fig 7A-D) and Müller glia extensions throughout the graft (Fig 7E).

**Figure 7:**
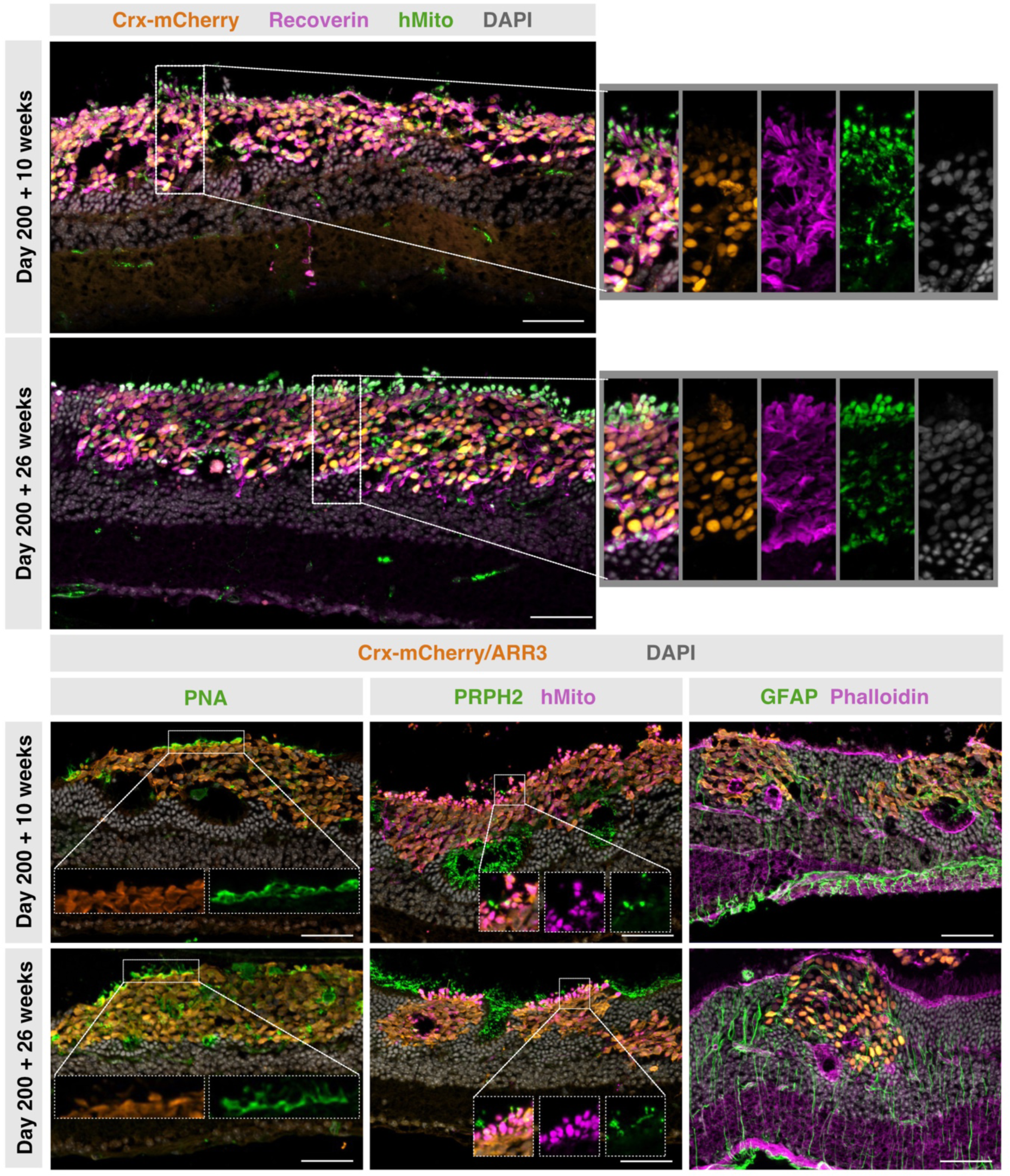
Crx-mCherry^+^ grafts also display extensive incorporation and polarisation – Retinal cryosections of Crx-mCherry^+^ grafts transplanted at D200 stained with recoverin, human mitochondria and DAPI shows (A) by 10 weeks large cell clusters incorporate into the host retina with areas of round mitochondria rich outgrowths towards the RPE and axon like extensions projected towards the inner nuclear layer (see zoomed area). (B) By 26 weeks grafts displayed even more abundant mitochondria rich outgrowths (see zoomed area). (C) PNA is bound in a more localised fashion towards the RPE. (D) Peripherin-2 is more extensively expressed at 26 weeks post transplantation and (D) Müller glia processes intermingle throughout the graft. Scale bars in all immunohistochemical images 50 μm

### Evidence for contact between host second order neurons and transplanted human cones

As cone axon-like protrusions were observed projecting towards the INL (Fig 2B,C), we aimed to assess whether there is also synaptic connectivity between transplanted photoreceptors 13 and host second order neurons in the highly interactive grafts. Immunohistochemical staining showed that both PKC*α*^+^ rod- and segretagogin^+^ cone-bipolar cell dendrites extended extensively into human cone clusters in areas where the donor cells are incorporated into the host ONL (Fig 8A; S3A). Further, calbindin^+^ horizontal cells also extended neural processes into the human cone grafts (Fig S3B). These observations indicate potential synaptic connections formed between donor cones and host second order neurons. To further investigate connectivity between donor and host cells, association between pre- and post-synaptic markers was assessed. As seen in Fig 8B, many examples of ribbon synapses labelled by CTBP2 within the graft can be found in close proximity to the bipolar cell post-synaptic marker mGluR6. This further supports putative synaptic connectivity between graft and host. Finally, the presence of typical photoreceptor ribbon synapses was confirmed by EM already at 10 weeks post-transplantation (Fig 8C).

**Figure 8:**
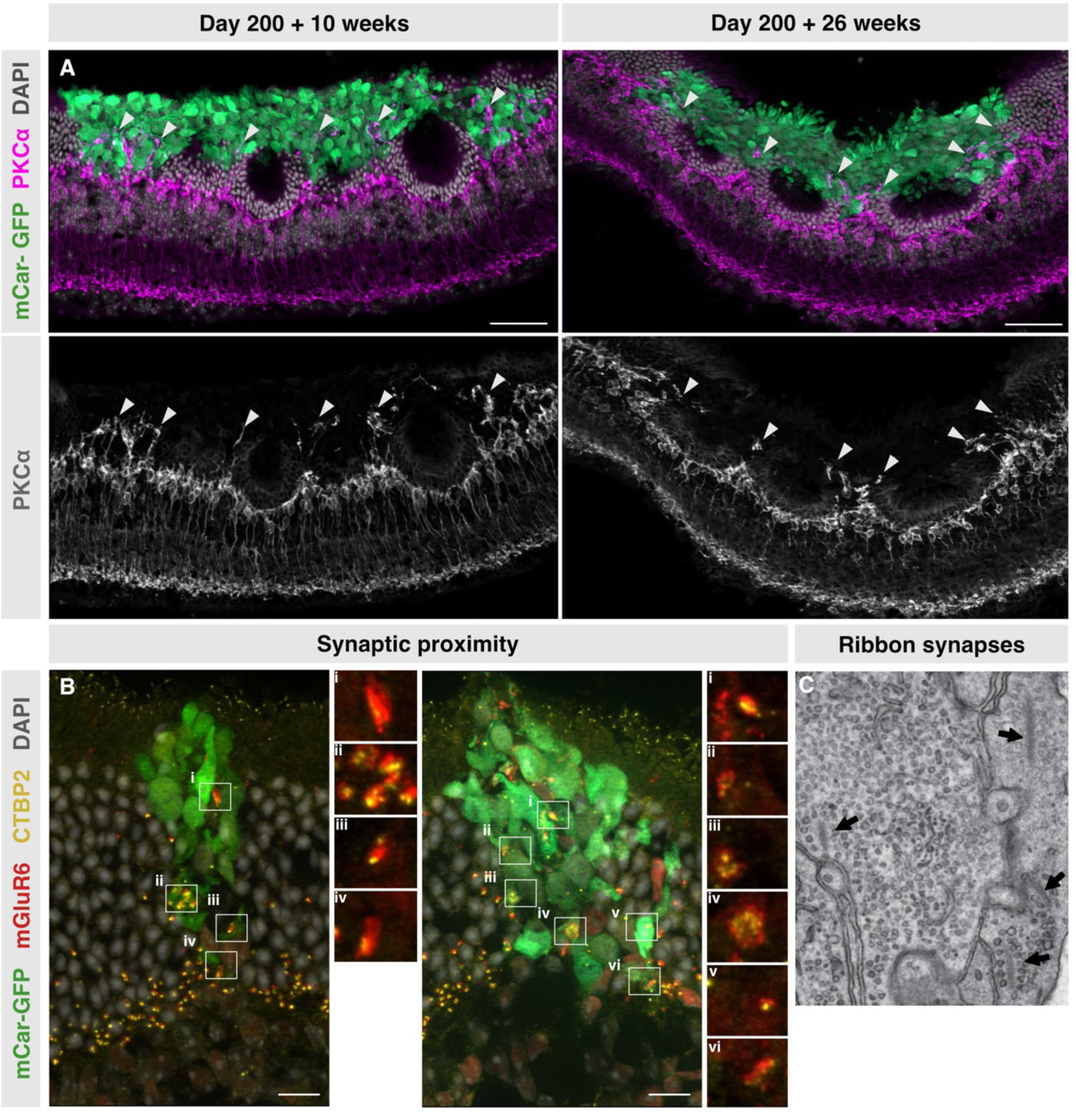
Putative synapse formation between transplanted human cones and host bipolar cells – Immunolabelled cryosections of Cpfl1 retina transplanted with mCar-GFP^+^ cells show (A) extensive dendrite extensions into the cone cell graft from PKCalpha^+^ rod bipolar cells. White arrowheads indicate areas of dendrite extensions. (B) Close association of the presynaptic ribbon synapse marker CTBP2 and the bipolar postsynaptic marker mGluR6. (C) Representative ribbons and vesicles, components of the photoreceptor presynapse, highlighted by arrows in a TEM image of an incorporated graft. Scale bars in all immunohistochemical images 50 μm and TEM 500 nm

To evaluate the functionality of these potential connections, we performed electrophysiological measurements using multi-electrode array (MEA) recordings. Here, due to technical challenges associated with cell mass localization of GFP causing severe bleaching, retinas containing Crx-mCherry cells were used. Robust and stable ON and OFF photopic light evoked responses (30 minutes of binary checkerboard white noise stimulation with stringent spike threshold settings to reduce artifacts) were detected in 5 of 9 transplanted eyes tested (Fig 9 A, B, E, F, G). However, low levels of photopic light responsiveness were also detected in non-transplanted regions of the same retina (Fig 9 E, F), but only following fluorescent stimulation, which was necessarily applied to locate the cell mass. Rods have been reported to respond to photopic light when over saturated (24). To eliminate potential endogenous oversaturated rod activity, the metabotropic glutamate receptor blocker L-AP4 was added during recording. L-AP4 blocks synaptic transmission between photoreceptors and all ON bipolar cells, including rod bipolar cells. Spike-triggered averaging was then used to categorise the ganglion cell response types (Fig 9 B, D). As expected, L-AP4 effectively quenched all ON RGC responses (Fig 9E, H). Moreover, OFF responses which are driven by cone bipolars remained only in the transplanted region (Fig 9F), strong evidence that the light-induced spiking activity is driven by the transplanted photoreceptors due to the lack of functional endogenous cones. Note that when the receptive field of the active ganglion cells was calculated, there was a high degree of overlap with the cell mass location (Fig 9C), further indicating that the transplant is driving the functional response to photopic light.

**Figure 9:**
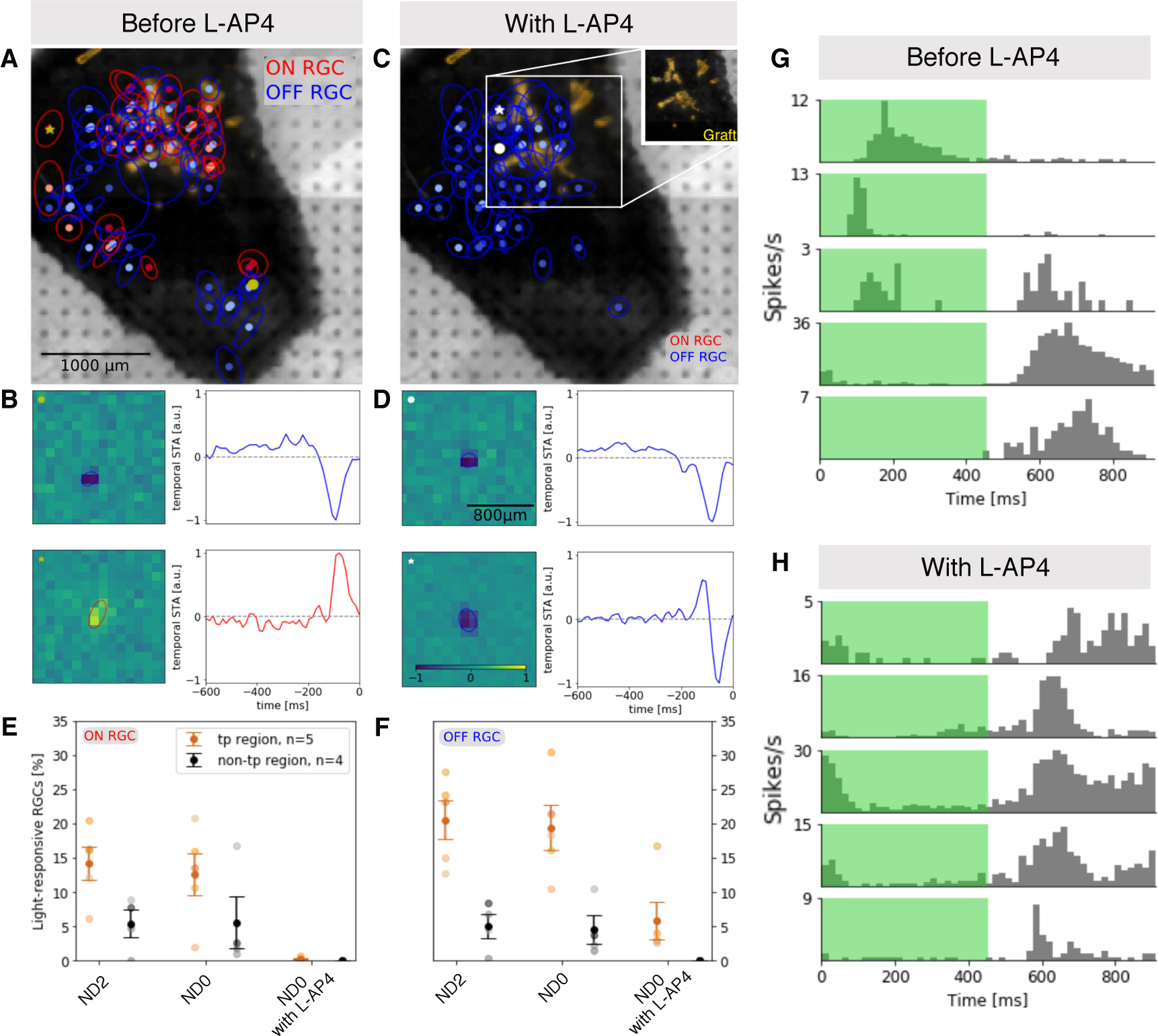
Increased RGC activity after Crx-mCherry^+^ photoreceptor transplantation – (A) Receptive Fields for ON and OFF RGCs detected following photopic stimulation (ND0). (B) Exemplary receptive fields and temporal STA for the two cells labeled with ∗ and ◦ in (A). (C) Receptive Fields where only OFF RGCs remain following photopic stimulation after addition of L-AP4 (ND0 after L-AP4). (D) Exemplary receptive fields and temporal STA for the two cells labeled with ∗ and ◦ in (C). Percentage of light responsive (E) ON RGCs and (F) OFF RGCs detected under mesopic (ND2), photopic (ND0) and photopic stimulation with the addition of L-AP4 (ND0 after L-AP4). (G-H) Response of 5 different RGCs during full-field photopic (ND0) ON-OFF flicker stimulation. The bin width is 20 ms and in total a number of 120 stimulus repetitions were performed. (G) Distinct ON, OFF and ON-OFF RGC responses to flicker stimulation before addition of L-AP4. (H) Distinct OFF RGC responses to flicker stimulation with addition of L-AP4 only remain in regions with transplant. RGC: Retinal ganglion cell, L-AP4: L-2-amino-4-phosphonobutyric acid.

## Discussion

In this study a human cone-specific GFP-reporter iPSC line facilitated the efficient enrichment of human cone photoreceptors from retinal organoids. The use of a local immune suppressant, monthly vitreal injection of triamcinolone acetonide, prevented the rejection of these human cells when transplanted into the Cpfl1 mouse subretinal space. This allowed long-term follow up over a 6-month period (26 weeks). With longer transplantation times, grafts interacted extensively with the host retina. These findings were confirmed through transplantations of a second photoreceptor-specific reporter iPSC line. Rather than forming a glial barrier, Müller glia intermingled throughout the graft, leading to the establishment of a common OLM between mouse and human cells. Second order neurons extended dendrites into the transplant forming potential synaptic connections. The incorporation of transplanted human cones into the host retina was accompanied by an improvement in cell polarisation and maturation of photoreceptor-specific morphological features, namely inner and outer 15 segments. Light detecting capacity and putative synaptic connectivity of transplanted human photoreceptors was further supported by light-evoked electrophysiological recordings of downstream retinal ganglion cells.

While human photoreceptor and rod specific ESC/iPSC reporter lines have been previously generated (23, 25–28), no human cone reporter PSC line has thus far been described. Based on immunohistochemical and transcriptional assessment, the herein presented mCar-GFP iPSC reporter line appears to robustly and specifically label human cone photoreceptors. This is not only useful for transplantation studies, but may also be of interest in other applications, e.g. studying human cone development or in the identification of human cone-specific cell surface markers. A previous study used viral labelling of L/M opsin cones to allow identification of cone cell surface markers. Not only does this exclude S-cones, but also, due to viral transduction efficiency, only around half of the total cone population was labelled (4). The resulting marker panel led to a maximal enrichment of ∼50% cones. A pan-cone reporter line would be of use in this context, as identification of cell surface markers is highly advantageous in a clinical setting where reporter or virally labelled fluorescent cells cannot be used.

In this study we show extensive incorporation of human cones and Crx-mCherry^+^ photoreceptors into the mouse ONL. This is to our knowledge the first report of such extensive incorporation of donor photoreceptors into the host retina from any species. Mouse into mouse photoreceptor transplantations largely result in material transfer rather than structural integration (18-20) – a mechanism that was ruled out in this study. As most recent studies of human photoreceptor suspension transplantation were either performed over a shorter time period and/or focused on transplantation into a fully degenerated retina (10–12, 21, 23), those experiments would not be expected to result in the aforementioned incorporation due to insufficient time (at 3 weeks only limited interaction was seen) or lack of ONL in which to incorporate. While three weeks after transplantation donor grafts mainly remain in the host subretinal space with only few contact points to the host ONL, at 10 weeks post-transplantation extensive incorporation was evident. However, areas where some host photoreceptors remained below the graft formed rosette-like structures reminiscent of outer retinal tubulations. Such tubulations are a well-known pathology upon retinal degeneration or damage (29) and it is assumed that this arrangement has a beneficial effect on photoreceptor survival when the RPE is defective. In the present case, rosette formation might thus be a response to the inaccessibility of RPE support in cases when the graft is sitting in between ONL and RPE. With longer times post-transplantation, and particularly with smaller clusters, human cones often fully incorporated into the host ONL, seeming to replace stretches of host photoreceptors with no obvious physical impediment to the host INL.

Single cell suspension studies have often been criticized for the lack of structure of the resulting graft (30). While in theory retinal sheet transplantation could provide pre-established structure, currently described studies suffer from extensive rosette formation and self-synapsing to graft second order neurons (5, 7–9, 31, 32). Sheet transplantations are surgically more challenging, particularly in the context of degenerative retinas where rupture of the tissue remains a potential risk. In this study, however, pre-purification of the transplanted cells was possible due to our photoreceptor-specific reporter lines and the used suspension technique. Unlike in other studies, the incorporated cones and Crx-mCherry^+^ photoreceptors appeared to become well polarized, with axon projections towards the INL and inner and outer segments towards the RPE. As photoreceptor loss is not complete until very late stages of blinding diseases, the remaining ONL may, as in this study, provide a structural framework for more organized integration of transplanted photoreceptors. This structural organization is likely aided by the close interaction with host Müller glia cells.

In the present work, graft maturation capacity was only observed upon incorporation into the host ONL. Through recovery of transplanted cells for next generation sequencing – a technique which has not yet been applied to photoreceptor transplants – we could show that in vivo matured cones from timepoints with extensive incorporation exhibit significant upregulation of visual transduction and outer segment related genes. With longer post-transplantation time, in vivo matured cones also increasingly expressed mitochondrial associated genes, which is noteworthy as mature cones are known to have very high energy requirements (33). Graft incorporation and maturation was further accompanied by close interaction with Müller glia, which not only intermingled throughout inner segment developing clusters, but even appeared to form a common OLM between human and mouse photoreceptor areas. Whether the Müller glia directly enhance maturation of the transplant remains to be proven, however, it is well known that glia are important supporters of neuronal function. Müller glia are critical for photoreceptor neurite outgrowth in both 2D and 3D culture systems (34, 35). Interestingly, Müller cells are also reported to play a role in organised outer segment assembly (36). In the present study, the outer segments that developed within older (D250) cone grafts, which incorporated to a much lesser extent and did not show much interaction with host Müller glia processes, were found to be highly disorganized.

While several studies have shown evidence of nascent outer segment formation – often in the organoids pre-transplantation rather than in the graft itself – these are usually small and/or with limited and disorganised discs (21, 37–41). A recent exception to this is the small but organized OS described by Ribeiro and colleagues, however no connecting cilium was shown (10). The outer segments seen in our study (D200 + 26 weeks) were not only tightly stacked and relatively well organized but were also seen sometimes to project from the inner segment via a connecting cilium, a feature which, to our knowledge, has not previously been reported in human photoreceptor suspension transplantations. Of note, organised outer segment formation including connecting cilium has been described in retinal sheet transplants (32), however these formed primarily within rosettes which would likely negatively affect function. A recent paper transplanted optogenetically engineered photoreceptors to circumvent the necessity for OS formation (42). While restored visual function was observed, this is limited to the specific wavelength of the optogene and has different kinetics to normal visual perception. Greater understanding and ideally modification of the factors required to encourage transplanted photoreceptors to develop and correctly form distinctive OS structures critical for light detection is of great importance if photoreceptor cell replacement therapy is to be an effective treatment modality.

Further interaction of host and donor tissue was seen at the level of the second order neurons. Rod and cone bipolar cells as well as horizontal cells extended dendrites into the transplant. Close proximity of pre- and postsynaptic ribbon synapse proteins supports the putative formation of synaptic connections. Of note, the putative synaptic connectivity occurred already at 10 weeks, preceding the extensive maturation of donor cells, as is also seen during development. Similar plasticity in second order neurons was already described in rd1 mice upon photoreceptor transplantation (10), but it is interesting that this effect is also seen in the Cpfl1 host where rod photoreceptors largely remain. This implies that the incorporated cell mass can replace existing connections, as dendritic remodeling of host second order neurons was observed only in areas of human cone incorporation. Similarly, in the aforementioned study, photopic light evoked responses by MEA were also reported (10). In our context, MEA recordings were complicated by the oversaturation of endogenous rods due to fluorescent cell mass localization, leading to a low level of background photopic response. For future studies, the injected cell number may be increased to expand graft area removing the need to locate by fluorescence, as per Ribeiro et al, where the transplantation of 500,000 donor cells not only increased graft area but also improved maturation compared to their previous studies using 150,000 cells (10). Regardless, using just 150,000 donor photoreceptors, we observed 3 to 4-fold higher proportion of both ON and OFF responsive RGCs under mesopic and photopic conditions when comparing regions containing transplanted cells with non-transplant containing regions. This is a strong indicator that the increased response is due to light-evoked responses transmitted from the graft. While the ON RGC contribution of the graft vs endogenous rods cannot be resolved definitively, the introduction of L-AP4 isolates cone OFF bipolar responses. As cones are dysfunctional or absent in the Cpfl1 host, any cone-OFF bipolar response should be due to newly formed connections to the graft. Indeed, all ON responses were quenched by L-AP4 and OFF responses remained only in the transplanted region, giving strong evidence that there is photopic light evoked signal transduction of transplanted cells through the cone OFF pathway. This indicates that the well matured and structurally incorporated photoreceptors in this study are functionally integrated and synaptically connected to the host retina.

In this study we describe the first human cone-specific reporter iPSC-line for the use of retinal organoid generation. Transplanted human cones and CRX^+^ photoreceptors extensively incorporated into a mouse model of cone degeneration. Incorporated grafts were well polarized and developed inner and outer segments. Further studies will be required to investigate details of the cellular and molecular requirements for structural incorporation and interaction with the host tissue allowing subsequent donor photoreceptor maturation. Such knowledge will be helpful to further optimize graft organization, OS formation and synaptic connectivity with the ultimate goal of improving visual perception. Nonetheless, the observed structural incorporation and subsequent in vivo polarisation and maturation of the human photoreceptors, second order neuron plasticity and the lack of physical impediment to synaptic connectivity is encouraging evidence that transplanted human photoreceptors may have the potential to integrate into the remaining outer nuclear layer of patients.

## Acknowledgements

The authors would like to thank the Stem Cell Engineering, Flow Cytometry, Light Microscopy and Deep Sequencing core facilities at the CMCB, Technische Universität Dresden for assistance in iPSC cell culture, cell sorting, microscopy and sequencing/ bioinformatic analysis, respectively. Additional technical assistance was provided by Jochen Hentschel, Sneha Prabhakara Shastry, Klara Schmidtke and Lynn Ebner.

This work was supported by the Bundesministerium für Bildung und Forschung (BMBF): ReSight - 01EK1613A to M.A., 01EK1613E to M.R. and G.Z)., and Deutsche Forschungsgemeinschaft (DFG): within the SPP2127 Program (AD375/7-1 to M.A.), and FZT 111 and EXC68. Supported by the Funding Programs for DZNE Helmholtz (M.K.); TU Dresden CRTD (M.K.); HGF ExNet-007 (M.K.); Bundesministerium für Bildung und Forschung (BMBF) ReSight (01EK1613A); DFG KA2794/5-1 SPP2127 (M.K.). This work received financial support from the State Ministry of Baden-Wuerttemberg for Economic Affairs, Labour and Tourism (M.R. and G.Z).V.B. acknowledges funding by the European Research Council (ERC-2020-PoC-966709 - iPhotoreceptors), by the Deutsche Forschungsgemeinschaft (SPP2127, EXC-2151-390873048 – Cluster of Excellence – ImmunoSensation2 at the University of Bonn) and the Volkswagen Foundation (Freigeist - A110720).T.K. and the EMF of the CMCB are supported by EFRE.

## Author contribution

S.G., K.T., and M.A. conceived this study. S.G., K.T., M.R., M.C., O.B., S.W., A.K., M.Z., M.V., T.K., O.G., M.K., V.B., G.Z. and M.A. designed and/or performed the experiments. S.G., K.T. and M.A. wrote this paper with input from all authors.

## Methods

### Vector production

The piggyBac vector backbone PB-TRE-dCas9-VPR (43) was a gift from George Church (Addgene plasmid, 63800). All promoter elements and open reading frames between the core insulator at the 5’ and the SV40 polA at the 3’ ends were removed using restriction enzymes and replaced with either PCR-amplified rod or cone reporter cassette. PCR products were introduced into the piggyBac vector backbone using Gibson assembly cloning (44). For the cone reporter cassette production, a PCR-amplified mouse cone arrestin promoter (mCAR) from LV-mCAR-eNpHR-EYFP (45) (gift from Botond Roska) was assembled with an EGFP followed by a downstream WPRE-BGH-pA element. Finally, a PCR-amplified ubiquitin C promoter (UBC)-blasticidin (Bla) cassette from vector pLV-TRET-hNgn1-UBC-Bla (46)(gift from Ron Weiss, Addgene plasmid, 61473) was further added to both vector assemblies resulting in reporter plasmids PB-hRHO-DsRed-WPRE-BGH-pA-UBC-Bla and PB-mCAR-EGFP-WPRE-BGH-pA-UBC-Bla. The plasmid DNA was transformed in chemically competent bacteria (One Shot® Stbl3™, Thermo Fisher Scientific) following the manufactureŕs protocol. The correct sequences were confirmed with Sanger sequencing. While RFP was also introduced under the Rhodopsin promoter, almost no RFP signal was detected even after 270 days in culture, however for the purposes of a cone transplantation study this was deemed irrelevant (data not shown).

### Generation of a hiPSC cone reporter line

The Personal Genome Project hiPS cell line PGP1 (47) was a gift from George Church (https://www.encodeproject.org, accession number: ENCBS368AAA). The cells were cultured on Matrigel coated wells (Corning, 354277) in mTeSR™1 medium (StemCell Technologies, 85850) and passaged in the presence of ROCK Inhibitor InSolution™ Y-27632 (Merck Millipore, 688001). The 4D-Nucleofector™ System (Lonza) was used to electroporate piggyBac and transposase vectors into PGP1 cells in suspension (X-Unit, P3 Primary Cell 4D-Nucleofector® X Kit L, program CB-156) following the manufacturer’s protocol. After nucleofections, cells were selected with 20 µg/ml Bla (Thermo Fisher Scientific, A1113903) for five days. The selected cells were seeded at low densities and propagated until each single cell formed a colony. Colonies were picked and genotyped using primers specifically binding to rod and cone reporter cassettes. The monoclonal cell line carrying both reporter cassettes (PGP1dR) at passage number 33-38 was used for all further experiments.

hRHO_for - GGATACGGGGAAAAGGCCTCCACGGCCACTAGTAGTTAATGATTAACCCG hRHO_rev - GACGTCCTCGGAGGAGGCCATGGTGGCTGCAGAATTCAGGGGATGACTCT

mCAR_for -

CTGGGGGGATACGGGGAAAAGGCCTCCACGGCCACTAGTGGTTCTTCCCATTTTGGCTAC

mCAR_rev -

GAACAGCTCCTCGCCCTTGCTCACCATGGTGGCTCTAGACCTCCAGCTCTGGTTGCTAAGCTGGC

### hiPSC maintenance and differentiation of retinal organoids

The mCar-GFP and Crx-mCherry iPSC lines (kind gift from Olivier Goureau – see (23)) were maintained in mTeSR1 (Stem cell technologies) on matrigel coated plates and split using ReleSR at room temperature (Stem cell technologies). Stem cells were differentiated to retinal organoids using an optimized protocol as previously described see also supplementary methods (17).

### FAC-sorting of reporter positive cells

Retinal organoids were dissociated in 20 U/ml papain, followed by gentle titration with a fire polished glass pipette and further processing as per the manufacturer’s instructions - Papain Dissociation System (Worthington). The cell pellet was resuspended in MACS buffer (0.5% BSA, 2 mM EDTA in PBS) to a concentration of ∼5 million cells per mL. The cell suspension was filtered through a 35 µm mesh and kept on ice for FAC sorting. An AriaII or AriaIII sorter was used to sort GFP^+^ or mCherry^+^ cells. Briefly, forward (FSC-A) and side scatter area (SSC-A) was used to discriminate cells from debris. Doublets were removed by gating FSC area vs height and by SSC height vs width. Dead cells were gated out using DAPI staining. Finally, GFP^+^ or mCherry^+^ cells were discriminated from auto fluorescent cells using GFP vs PE or APC.

### Animals

Adult cone photoreceptor function loss 1 (Cpfl1) mutant (7-14 week-old) were used as recipients for cell transplantation. Mice were maintained in a 12-hour Light/Dark cycle with ad libitum access to food and water.

### Transplantations

Following FAC sorting, GFP^+^ or mCherry^+^ cells were resuspended in MACS buffer (150 000 cells/µl) and injected into the subretinal space of host eyes as previously described -see also supplementary (48). Directly following cell transplantation, 1 µL of preservative free triamcinolone acetonide suspension (80 µg /µL in NaCl prepared by University Clinic Pharmacy, Dresden) was injected into the vitreous using a hand held 10 µl Hamilton syringe with a blunt 34-Gauge needle. Triamcinolone vitreal injections were repeated on a monthly basis.

### Immunohistochemistry

Immunohistochemistry was performed as described previously (18) see supplementary methods for details. For immunocytochemistry following dissociation and sorting of the retinal organoid cells, cells were resuspended in RM2 media and laminin was added to each fraction. From each fraction 50 000 cells were plated into flexiperm wells on a PDL coated slide. Cells were incubated at 37°C for 2 hours to allow attachment. Cells were then fixed for 15 minutes at room temperature (RT), washed 3 times with PBS and stained as per frozen sections above. Frozen sections and plated cells were mounted following antibody staining using AquaPolymount (Polysciences, Heidelberg, Germany) and imaged using a Zeiss Apotome ImagerZ2 (Zeiss, Heidelberg, Germany).

### Transmission Electron Microscopy (TEM) and Correlative Light Electron Microscopy (CLEM)

TEM of transplanted cones was performed as previously described (49, 50). CLEM of immunolabeled sections was performed as described previously (51, 52) TEM imaging was performed with a Jeol JEM1400 Plus transmission electron microscope (camera: Ruby, Jeol) running at 80kV acceleration voltage.

### SmartSeq2

Whole eye cups of transplanted eyes or organoids maintained in culture from the same differentiation round were dissociated with papain as described above (Papain dissociation kit 20U). Cell were resuspended in MACS buffer and filtered through a 35 µm mesh before FAC-sorting and sequencing - method was modified based on (53) see supplementary for details.

### Transcriptomic Analysis

Basic quality control of the resulting sequence data was done with FastQC (v0.11.6) (https://www.bioinformatics.babraham.ac.uk/projects/fastqc/) and the degree of mouse contamination was assessed with FastQ-Screen (v0.9.3) (https://www.bioinformatics.babraham.ac.uk/projects/fastq_screen). Reads originating from mouse were removed with xengsort (v2021-05-27)(54). Reads were aligned to the human reference genome hg38 using the aligner gsnap (v2020-12-16)(55) with Ensembl 92 human splice sites as support. Uniquely mapped reads were compared based on their overlap to Ensembl 92 human gene annotations using featureCounts (v2.0.1)(56) to create a table of fragments per human gene and sample. Normalization of raw fragments based on library size and testing for differential expression between the different cell types/treatments was performed with the R package DESeq2 (v1.30.1) (57). Sample to sample Euclidean distance, Pearson’ and Spearman correlation coefficient and principal component analysis based upon the top 500 genes with the highest variance were computed to explore correlation between biological replicates and different libraries. To identify differentially expressed genes, counts were fitted to the negative binomial distribution and genes were tested between conditions using the Wald test of DESeq2. The comparison of the GFP-positive vs the GFP-negative fraction included the age as covariate while all other comparisons just included the specific groups. Resulting p-values were corrected for multiple testing with the Independent Hypothesis Weighting package (IHW 1.18.0) (58, 59). Genes with a maximum of 5% false discovery rate (padj ≤ 0.05) were considered as significantly differentially expressed.

### Electrophysiological recordings with multi electrode array

A Glass MEA with 256 electrodes of 30 μm diameter and a spacing of 200 μm spanning an area of 3 mm × 3 mm (256MEA100/30iR-ITO) with recording headstage (MEA256-System, Multi Channel Systems MCS) below the microscope was used for all experiments.

The preparation of ex vivo retina was performed in carbonated (95% O2, 5% CO2) Ames’ medium (Ames A 1420, Sigma Aldrich + NaHCO_3_). Following the euthanasia of the mouse, the eyes were opened via a small needle incision above the ora serrata. After removal of the lens, the eye was cut in half and the graft located with stereomicroscope (Leica M80), equipped with a fluorescent illumination unit. The retina with the graft was then separated from the sclera and RPE, trimmed with a scalpel and the vitreous removed. The retina was placed ganglion cell side up on a filter paper and transferred retinal ganglion cell (RGC) side down on to the coated (Cell-Tak, Corning), as described in detail in a previous report (60) recording electrodes and filter paper removed. The other half of the retina was prepared in the same way as a reference sample.

A patterned light stimulus created by an oLED display (DSVGA monochrome green XLT, eMagin) in combination with the software GEARS (61) was used, allowing for binary checkerboard white noise (bwn) stimulation. The oLED is coupled to the microscope with an adapter and its light is projected onto the sample though a 2.5x objective. The oLEDs power was derived as P=0.7 μW for full-field illumination, which can be calculated into photoisomerizations equaling to approx. 1·10^5^ R*/photoreceptor/s for both rods and m cones. We generated pseudo-random, binary (green and black) checkerboard stimuli - where at every stimulus frame the intensity of each checker was drawn from a binary distribution - with temporal frequency of 38 Hz (frame duration of 26ms) and a total duration of 25 min, with a resolution of 30 pixel x 30 pixel, resulting in an illuminated area of 3.2 mm x 4.2 mm.

During RGC activity recording the MEA chamber was continuously perfused with Ames’ solution at a rate of 2-4 mL/min. The temperature of the MEA chamber was maintained at ∼ 36°C by heating the bottom of the recording chamber and the perfusion inlet. To assure RGC OFF responses are driven by injected photreceptors, experiments were performed before and after addition of the mGluR6 blocker L-AP4 (50 μM, Tocris Cat. No. 0103). Extracellular voltages were recorded using the software MCRack (MCS) and preprocessed using a 2nd order Butterworth highpass filter (300Hz), before spike detection -see supplementary for details.

### Image processing

Images and graphs were processed and generated using Image J (National Institutes of Health), Zen Blue Software (Zeiss), Affinity Designer (Serif Ltd), and graphpad Prism 7

Recoverin and cone arrestin quantification of ICC was performed using cell profiler 3.1.9. Total graft area was quantified using Zen Blue image analysis wizard, and then each individual GFP^+^ cell cluster with associated area was categorized manually as ONL contact – cluster made contact with the ONL but mostly remained in the subretinal space, partially incorporated – the cluster was in line with the host ONL, however gaps or rosette like structures from remaining host ONL reside below the graft area, or fully incorporated – the cluster replaced sections of host ONL without gaps or rosettes from the host.

Panther was used for gene enrichment analysis (62). Differentially expressed genes from our data set were run through the statistical overrepresentation test function using the whole human genome as reference list. Fisher’s exact test with calculated false discovery rate was selected and output was condensed by hierarchical clustering of GO-terms to reduce repetitive pathway findings. Morpheus (https://software.broadinstitute.org/morpheus) was used to create heat maps.

### Statistical analyses

Statistical significance was calculated using a one-way ANOVA with Tukeys multiple comparison tests. Statistical significance is represented in the figures as follows: *, *p* < .05; **, *p* < .01; ***, *p* < .001; ****, *p* < .0005, n.s.: not-statistically significant. Detailed statistical analysis of transcriptional data and spike sorting see respective sections.

### Study approval

All animal experiments were approved by the ethics committee of the TU Dresden and the Landesdirektion Dresden (approval number: TVV 16/2016 and TVV 38/2019). All regulations from European Union, German laws (Tierschutzgesetz), the ARVO statement for the Use of Animals in Ophthalmic and Vision Research and the NIH Guide for the care and use of laboratory work were strictly followed for all animal work.

