## Supplementary methods and figures for "Extensive incorporation, polarisation and improved maturation of transplanted human cones in a murine cone degeneration model"

### Supplementary Material

#### Transplantations

Mice were anesthetized via intraperitoneal injection of medetomidine hydrochloride (1 mg/kg, Domitor, Orion Pharma) and ketamine (0.75mg/10g body weight; Ratiopharm) and pupils dilated with 2.5% phenylephrine/0.5% tropicamide (TU Dresden Pharmacy). The mouse head was secured and positioned with a head holder (Leica, Wetzlar, Germany) so that the optic nerve was visible in the central fundus. A hole was made in the ora serrata with a sharp 30-gauge needle (VWR, Germany). 1 µL of cell suspension then 0.2 µL of air was drawn into a 5 µl Hamilton syringe with a blunt 34-Gauge needle. The syringe was attached to a micromanipulator and transvitreally inserted into the eye under visual control aided by a microscope. The needle was placed nasally in the subretinal space, the air bubble quickly injected to create a bleb and then the cell suspension slowly injected. Anesthesia was reversed by intraperitoneal injection of atipamezole (10mg/kg, Antisedan, Orion Pharma).

#### Differentiation of retinal organoids

Briefly, when hiPSC reached 60% or 80% confluence (mCar-GFP and Crx-mCherry respectively) these were detached to small cell clusters with ReLeSR and resuspended in Matrigel (growth-factor reduced, BD Biosciences). Matrigel was allowed to gel for 5-7min at room temperature and was then gently dispersed into small clumps in N2B27 media (1:1 DMEM/F12: Neurobasal A media, 1% B27+VitaminA, 0.5% N2, 1% penicillin/streptomycin, 1% GlutaMAX, 0.1 mM 2-mercaptoethanol). This suspension was added to 6-well ultra-low attachment plates (Nunclon Sphera, Thermo Fisher) allowing floating neuroepithelial cysts to form within the first few days of culture. Cysts were then plated on Matrigel coated plates on day 5, followed by gentle Dispase detachment (Stem cell technologies) on day 13. Detached clusters were transferred to 9 cm ultra low attachment plates (Nunclon Sphera, Thermo Fisher) in B27 media (DMEM/F12, 1% B27 without Vitamin A, 1% penicillin/streptomycin, 1% GlutaMAX, 1% NEAA, 0.1% Amphotericin B). On day 30 retinal epithelial domains were manually isolated using surgical tweezers (Fine Science Tools, Dumont No. 5). From D25, B27 media was supplemented with 10% FBS and from D100 N2+FBS media was used (DMEM/F12, 1% N2, 10% FBS, 1% penicillin/streptomycin, 1% GlutaMAX, 0.1% Amphotericin B). Synthetic retinoid analogue EC23 (0.3 µM) was supplemented from D25 to D120. Media was refreshed with 50% media change every 2-3 days from day 13 onwards.

### Immunohistochemistry

Experimental animals were sacrificed by cervical dislocation. Eyes were enucleated and fixed in 4% paraformaldehyde (PFA) solution for 1 hour at 4°C. The cornea and lens were then dissected away and the remaining eye cup cryopreserved in 30% (wt/vol) sucrose solution overnight at 4°C. Similarly, organoids were fixed in 4% PFA for 30 minutes at room temperature followed by 10% sucrose 1hr, 30% sucrose 3 hr and 50% sucrose overnight at 4°C. Eyes and organoids were embedded in NEG50 (Thermo Scientific, Schwerte, Germany), and cryo-sectioned to 12 µm thickness using a microtome (Micron HM560, Thermo Scientific). Frozen sections were air-dried briefly at room temperature (RT) and hydrated for 5 minutes in phosphate buffered saline (PBS) followed by blocking (0.3% Triton X-100 solution with 1% bovine serum albumin (BSA) and 5% donkey serum) for 1 hour at RT. Antigen retrieval was performed prior to blocking for CTBP2 and mGluR6 staining (1mM

EDTA, 0.05% Tween20, pH 8 for 30 min at 70°C). Sections were incubated overnight at 4°C with primary antibodies (see supplementary table 1) in blocking solution. Slides were washed with PBS three times to remove unbound primary antibody. Corresponding secondary antibodies coupled to the fluorescent dyes AlexaFluor 488, AlexaFluor 546, Cy2, Cy3, or Cy5, at a dilution of 1:1000, were incubated for 1 hour 30 minutes at RT together with 4',6-diamidino-2-phenylindole (DAPI; 1:15,000; Sigma).

| Antibody | Species | Dilution | Source | Catalogue no. |
|---|---|---|---|---|
| Calbindin | mouse | 1 :1000 | Swant | 300 |
| Cone arrestin - human sp. | mouse | 1:100 | Gift from Peter MacLeisch | |
| CRALBP | mouse | 1:100-1:200 | invitrogen | MA1-813 |
| CRX | rabbit | 1:500 | Gift from Elly Tanaka | |
| CTBP2 | mouse | 1 : 2500 | BD | 612044 |
| GFAP | rat | 1:500 | Millipore | 345860 |
| GFP | chicken | 1 : 500 | Abcam | ab13970 |
| mGluR6 | rabbit | 1:2000 | Alomone | AGC-026 |
| HuC/D biotin-conjugated | mouse | 1:100 | Invitrogen | A-21272 |
| Iba1 | rabbit | 1:500 | Wako | 019-19741 |
| human mitochondria | mouse | 1:500 | abcam | ab92824 |
| human mitochondria | mouse | 1:500 | abcam | ab3298 |
| Nrl | goat | 1:200 | R&D | AF2945 |
| L/M Opsin | Rabbit | 1:1000 | Millipore | AB5405 |
| S-opsin | goat | 1:200 | Santa Cruz | sc-14363 |
| Peripherin, PRPH2 | rabbit | 1:200 | Thermo fisher | 18109-1-AP |
| PKCα | rabbit | 1:300 | Santa Cruz | sc-208 |
| PNA biotinylated | | 1 : 1000 | Vector Laboratories | B-1075 |
| Recoverin | rabbit | 1:1500 | Millipore | AB5585 |
| Rhodopsin (opsin RET-P1) | mouse | 1:1000 | Sigma | O4886 |
| Secretagogin | sheep | 1:300 | Biovendor | RD184120100 |
| SOX2 | goat | 1:200 | santa cruz | sc-17320 |

### Transmission electron Microscopy

Retinae were fixed in 4% formaldehyde (prepared from paraformaldehyde prills) in 100mM phosphate buffer (PB) and the region of interest (ROI) with transplanted GFP-positive cells was cut out using the Leica MZ10F stereomicroscope equipped with epifluorescence. Dissected samples were postfixed in modified Karnovsky's fixative (2% glutaraldehyde, 2% paraformaldehyde in 50 mM HEPES) overnight at 4°C (1). Samples were washed and further postfixed in 2% aqueous $OsO_4$ solution containing 1.5% potassium ferrocyanide and 2mM $CaCl_2$. After washing, samples were incubated in 1% thiocarbohydrazide, washed again, and contrasted in 2% aqueous $OsO_4$ for a second time. After washing, samples were *en-bloc* contrasted with 1% uranyl acetate/water, washed again in water, dehydrated in a graded ethanol series and infiltrated in the epon substitute EMBed 812. After embedding, samples were cured at 65°C overnight. Ultrathin sections were cut with a Leica UC6 ultramicrotome and collected on formvar-coated slot grids. Sections were stained with lead citrate (2) and uranyl acetate.

### Correlative light and electron microscopy

ROIs from retinae fixed in 4% formaldehyde/100 mM PB were dissected under a fluorescence stereomicroscope (Leica MZ10F), dehydrated in a graded series of ethanol, infiltrated and embedded in Lowicryl K4M at progressively lower temperatures (PLT, down to -35°C, (3). Polymerization occurred by UV-irradiation at -35°C for 48hrs. Ultrathin sections were mounted to formvar-coated mesh grids, blocked with 1% bovine serum albumin in PBS, incubated with mouse anti-cone arrestin 3, a rabbit-anti-mouse bridging antibody, protein A 10 nm gold (immunogold), and goat-anti-rabbit Alexa488 (immunofluorescence). In addition, the labeled sections were treated with DAPI (4,6-diamino-2-phenyl-indole) to stain the nuclei for immunofluorescence microscopy. The labeled grids were mounted in glycerol-water (1:1) and analyzed with a Keyence Biozero 8000 fluorescence microscope. Regions with transplanted cones were identified by fluorescence and selected for further analysis at the TEM. Selected grids were demounted, washed in water, contrasted with uranyl acetate and dried for TEM inspection.

### SmartSeq2

30 $GFP^+$ cells were sorted via FACS into a 96 well plate or PCR strip tubes containing 2 µl of nuclease free water with 0.2% Triton-X 100 and 4 U murine RNase Inhibitor (NEB), spun down and frozen at -80°C. After thawing, 2 µl of a primer mix (5 mM dNTP (Invitrogen), 0.5 µM dT-primer*, 4 U RNase Inhibitor (NEB)) was added to the cell lysate. RNA was denatured for 3 minutes at 72°C and reverse transcribed at 42°C for 90 min after filling up to 10 µl with RT mix for a final concentration of 1x superscript II buffer (Invitrogen), 1 M betaine, 5 mM DTT, 6 mM $MgCl_2$, 1 µM TSO-primer*, 9 U RNase Inhibitor and 90 U Superscript II. After synthesis, the reverse transcriptase was inactivated at 70°C for 15 min. The cDNA was amplified using Kapa HiFi HotStart Readymix (Roche) at a final 1x concentration and 0.1 µM UP-primer* under following cycling conditions: initial denaturation at 98°C for 3 min, 18 cycles [98°C 20 sec, 67°C 15 sec, 72°C 6 min] and final elongation at 72°C for 5 min. The amplified cDNA was purified using 0.6x volume of Sera-Mag SpeedBeads (GE Healthcare) resuspended in a buffer consisting of 10 mM Tris, 20 mM EDTA, 18.5 % (w/v) PEG 8000 and 2 M sodium chloride solution. The cDNA was eluted in 12 µl nuclease free water and its concentration was measured with Infinite M $Nano^+$ (Tecan) in 384 well black flat bottom low volume plates (Corning) using AccuBlue Broad range chemistry (Biotium).

For library preparation 3 ng cDNA was tagmented using 0.5 µl TruePrep Tagment Enzyme V50 and 1x TruePrep Tagment Buffer L (TruePrep DNA Library Prep Kit V2 for Illumina, Vazyme), followed by an incubation step at 55 °C for 10 min. Next, Illumina indices were added during PCR (72 °C 3 min, 98 °C 30 sec, 12 cycles [98 °C 10 sec, 63 °C 20 sec, 72 °C 1 min], 72°C 5 min) with 1x concentrated KAPA Hifi HotStart Ready Mix and 300 nM dual indexing primers. After PCR, libraries were purified with 0.9x volume Sera-Mag SpeedBeads, followed by a size selection with 0.6x and 0.9x volume of beads to enrich fragments of a size between 200 and

800 bp. Sequencing was performed after quantification with a Fragment Analyzer, either on Illumina Nextseq500 or Novaseq 6000 with an average sequencing depth of 7 mio fragments.

dT-primer: C6-aminolinker-AAGCAGTGGTATCAACGCAGAGTCGAC TTTTTTTTTTTTTTTTTTTTTTTTTTTTTTVN, where N represents a random base and V any base beside thymidine;
TSO-primer: AAGCAGTGGTATCAACGCAGAGTACATrGrGrG, where rG stands for ribo-guanosine;
UP-primer: AAGCAGTGGTATCAACGCAGAGT

### MEA Spike Triggered Average and Receptive Field calculations

For spike sorting, we first performed threshold crossing to detect all spike events. The detection threshold (th) was chosen by manually checking the performance of the sorting:

$$\text{th} = 8 \cdot \text{median}(\text{dev}/0.6745), \text{ with dev} = \sqrt{\sum_i (X_i - \bar{X})^2}. \qquad (1)$$

$X = (X_1, X_2, ..., X_n)$ is taken from the first 10 seconds of the spontaneously recorded activity. The high threshold was chosen to reduce artifacts and focus on close-by cells to the electrodes (stronger waveform). After detecting the spike events, we clustered them using principal component analysis (PCA) based on 3 eigenvectors into a maximum of 6 clusters based on the KlustaKwik algorithm (4). The detected clusters represent single cell activities.

Temporal resolution, an important factor determining success in vision restoration, can be assessed by evaluating the RGC response to bwn stimulation. Averaging over the spike-triggered stimulus ensemble (STE), which is the collection of all stimuli that were followed by a spike within a certain time window, leads to the spike-triggered average (STA)(5).

$$k(\tau_i) = \frac{1}{n} \sum_{k=1}^{n} s(t_k - \tau_i), \qquad (2)$$

with n the number of spikes and $\tau_i$ the time shift relative to the spike. The cells STA is given by

$$k = (k(\tau_{max}), ..., k(\tau_0)). \quad (3)$$

To decide if a detected STA is resulting from noisy cell response or robust cell response, we estimated the extrema threshold ratio as $th_{ratio} = 7\sigma$.

The STA can be derived on a pure temporal basis or on a spatial and temporal basis. If bwn stimulation is performed both temporal and spatial, spatial receptive fields of responsive cells can be estimated.

To determine the spatial extent of the Gaussian shaped STA, a 2D Gaussian function was fitted as follows:

$$\begin{aligned} a &= \frac{\cos(\theta)^2}{2\sigma_x^2} + \frac{\sin(\theta)^2}{2\sigma_y^2} \quad (4) \\ b &= -\frac{\sin(2\theta)}{4\sigma_x^2} + \frac{\sin(2\theta)}{4\sigma_y^2} \end{aligned}$$

$$\begin{aligned} c &= \frac{\sin(\theta)^2}{2\sigma_x^2} + \frac{\cos(\theta)^2}{2\sigma_y^2} \\ g(x,y) &= O + A \cdot \exp\left(-\left[a \cdot (x-x_0)^2 + 2b \cdot (x-x_0) \cdot (y-y_0) \right.\right. \\ &\quad \left.\left. + c \cdot (y-y_0)^2\right]\right), \end{aligned}$$

with offset O, peak height A, center of the ellipsoid ($x_0$, $y_0$) and angle θ. Koehler et. al (Koehler, Akimov, and Renteria 2011) use a two-dimensional Gaussian fit to the peak frame of the STA (i.e., the frame containing the largest deviation from the mean) as a spatial representation of the receptive field. Here, to reduce influence of noisy pixel and to highlight the STA - cell center pixel, the recorded STA images for time points between 80 – 24 ms (binned in 8 ms) pre spike were each normalized to the respective extrema value and then averaged.

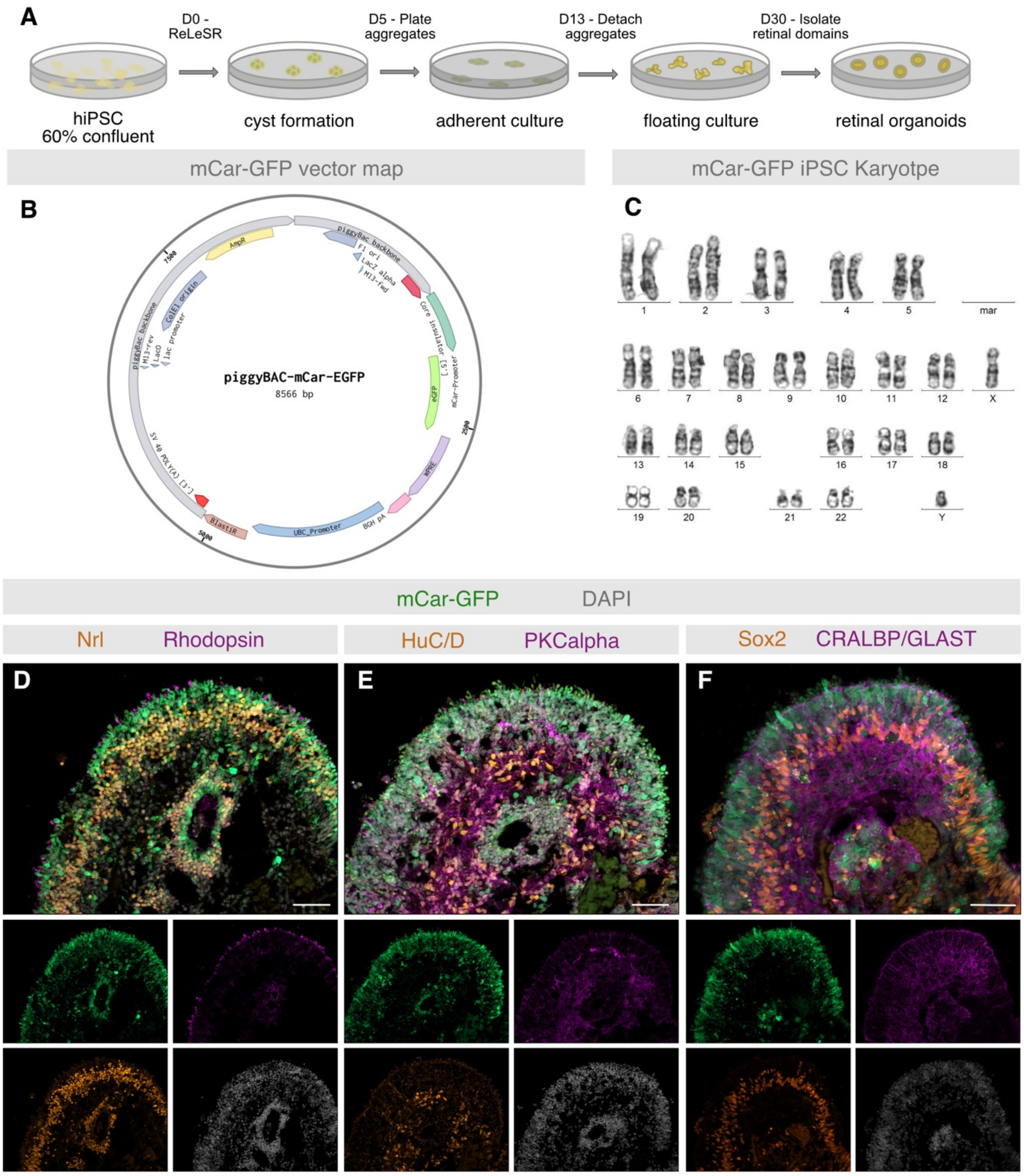


Supplementary Figure 1: Generation and validation of a cone iPSC cell line – (A) Schematic of retinal organoid generation. (B) mCar-GFP vector map. (C) Karyotype of the resulting mCar-GFP iPSC line. Immunostaining of mCar-GFP retinal organoid cryosections show little overlap of GFP with (D) rod markers Nrl and rhodopsin, (E) retinal ganglion cell and amacrine marker HuC/D and bipolar cell marker PKCalpha, (F) Müller glia markers SOX2, CRALBP and GLAST. Scale bars in all immunohistochemical images 50 µm.

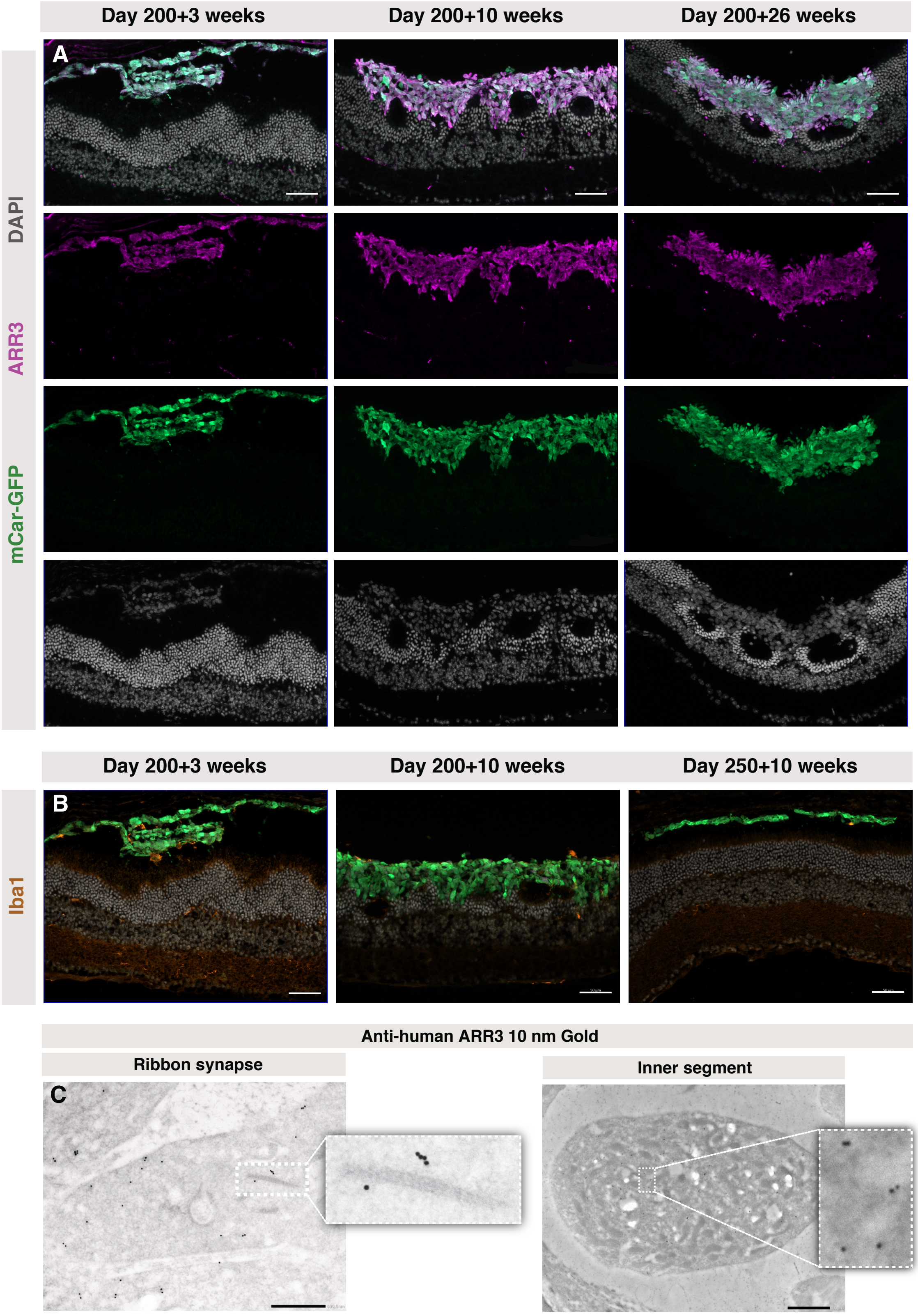


Supplementary Figure 2: Grafts express ARR3 and cause minimal immune reaction – Immunolabelled cryosections of Cpfl1 retina transplanted with mCar-GFP cells (A) express human ARR3 and mCar-GFP across all timepoints. DAPI staining shows that human cones have much larger, less dense nuclei than the remaining mouse host photoreceptors. (B) Few Iba1 positive cells are identified in the graft area. (C) Immunogold labelling with anti-human ARR3 which was detected with protein A 10nm gold particles shows photoreceptor specific features of ribbon synapse and inner segments originating from human cone cells transplanted into the mouse. Scale bars in all immunohistochemical images 50 µm and for TEM images 1 µm

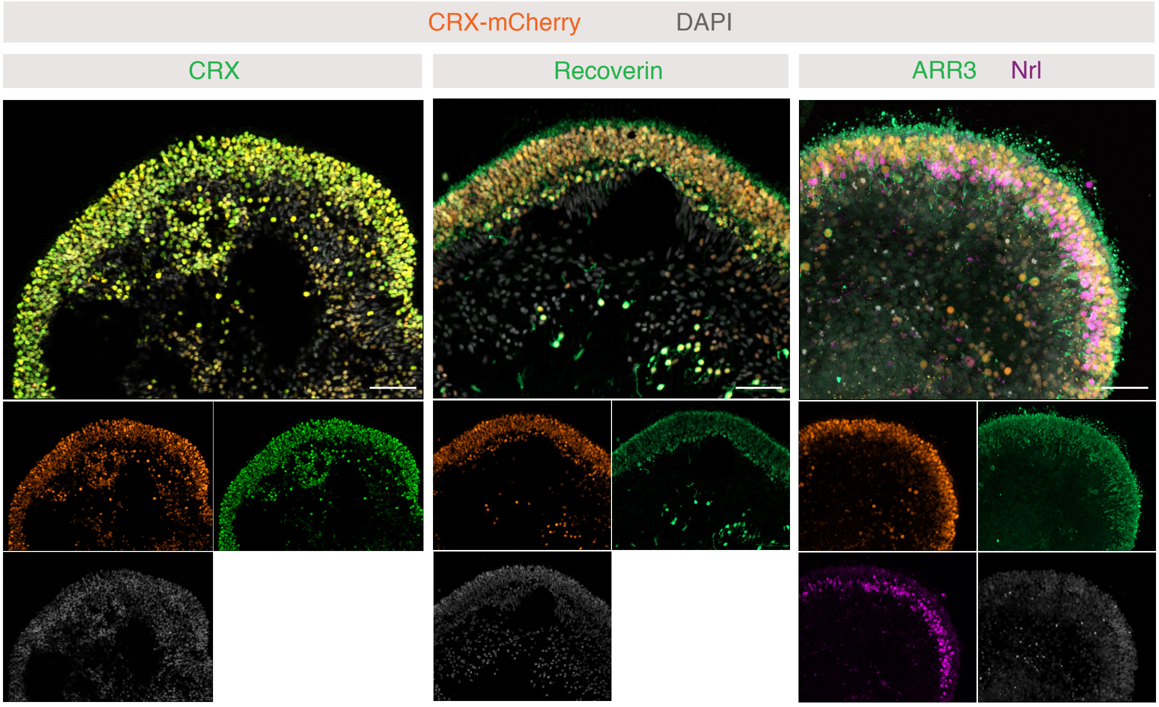


Supplementary Figure 3: Crx driven mCherry labels rods and cones – Immunostaining of Crx-mCherry retinal organoid cryosections show mCherry overlaps with Crx, Recoverin, ARR3 and Nrl. Scale bars in all immunohistochemical images 50 µm

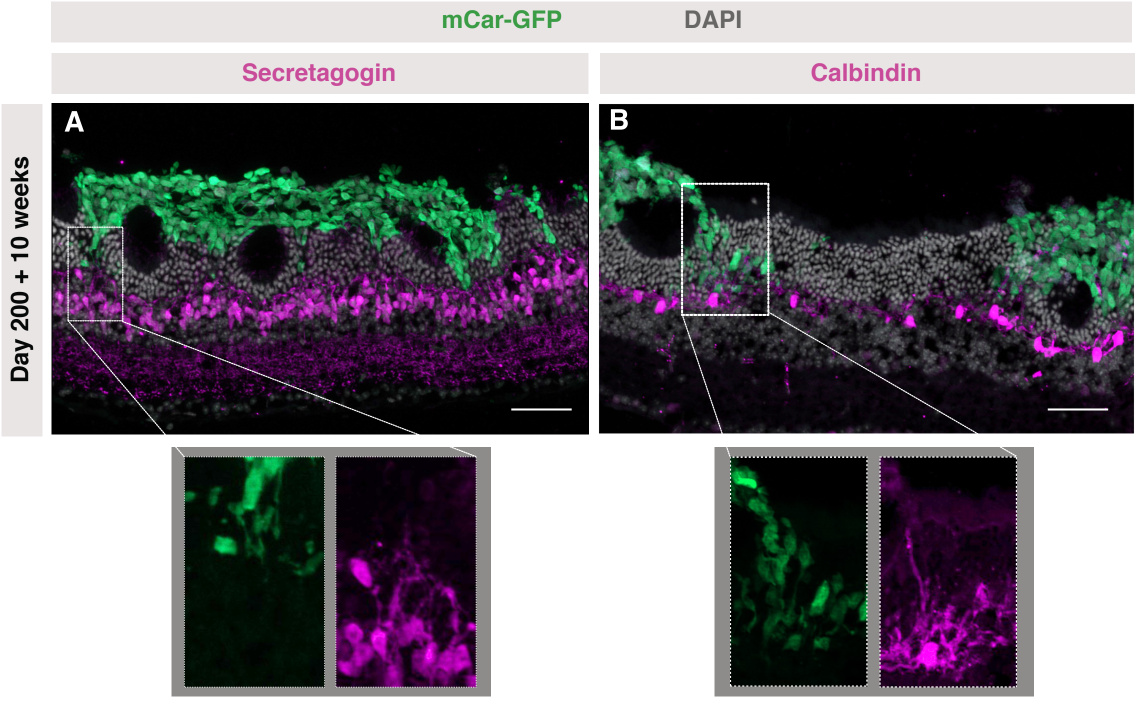


Supplementary Figure 4: Cone bipolar and horizontal cells extend dendrites into the graft – Immunolabelled cryosections of Cpfl1 retina transplanted with mCar-GFP cells show dendrite extensions into the cone cell graft from (A) secretagogin+ cone bipolar cells and (B) calbindin+ horizontal cells. Scale bars in all immunohistochemical images 50 µm
